# Stressor-Selective Sympathetic Preganglionic Modules for Organ-Biased Control

**DOI:** 10.64898/2026.09.07.749966

**Authors:** Serika Yamada, Shuntaro Uchida, Masafumi Tsurutani, Dooseon Cho, Mitsutaka Kadota, Takefumi Kondo, Kazunari Miyamichi, Haruaki Sato

## Abstract

The sympathetic nervous system coordinates organ function during stress, yet the spinal organization that converts autonomic commands into selective peripheral outputs remains poorly understood. Here, we combined spatial transcriptomics, immediate early gene mapping, anatomical tracing, and functional perturbation to define the cellular logic of spinal sympathetic preganglionic neurons (SPNs) in mice. SPN transcriptomic subtypes were spatially organized by spinal segment and sex and recruited in distinct combinations by physiological stressors, revealing stressor-selective sympathetic output modules. Focusing on cold-responsive *neurotensin*-expressing (*Nts*+) SPNs in the lower thoracic spinal cord, we found that they formed a distinct output channel to the lower sympathetic trunk and aorticorenal ganglia, promoted female-biased mobilization of white adipose tissue lipids, and were required for cold tolerance when food was unavailable. These findings establish a cell-type-resolved spinal architecture that links physiological demand to organ-biased sympathetic output and identifies the preganglionic layer as a key organizer of brain–body control.

**Highlights:**

- SPN subtypes are spatially organized by spinal segment and sex
- Distinct stressors selectively recruit distinct sets of SPN subtypes
- Cold-responsive *Nts*+ SPNs form a distinct sympathetic output module
- *Nts+* SPNs drive female-biased fat mobilization and cold tolerance

**Graphical Abstract:** 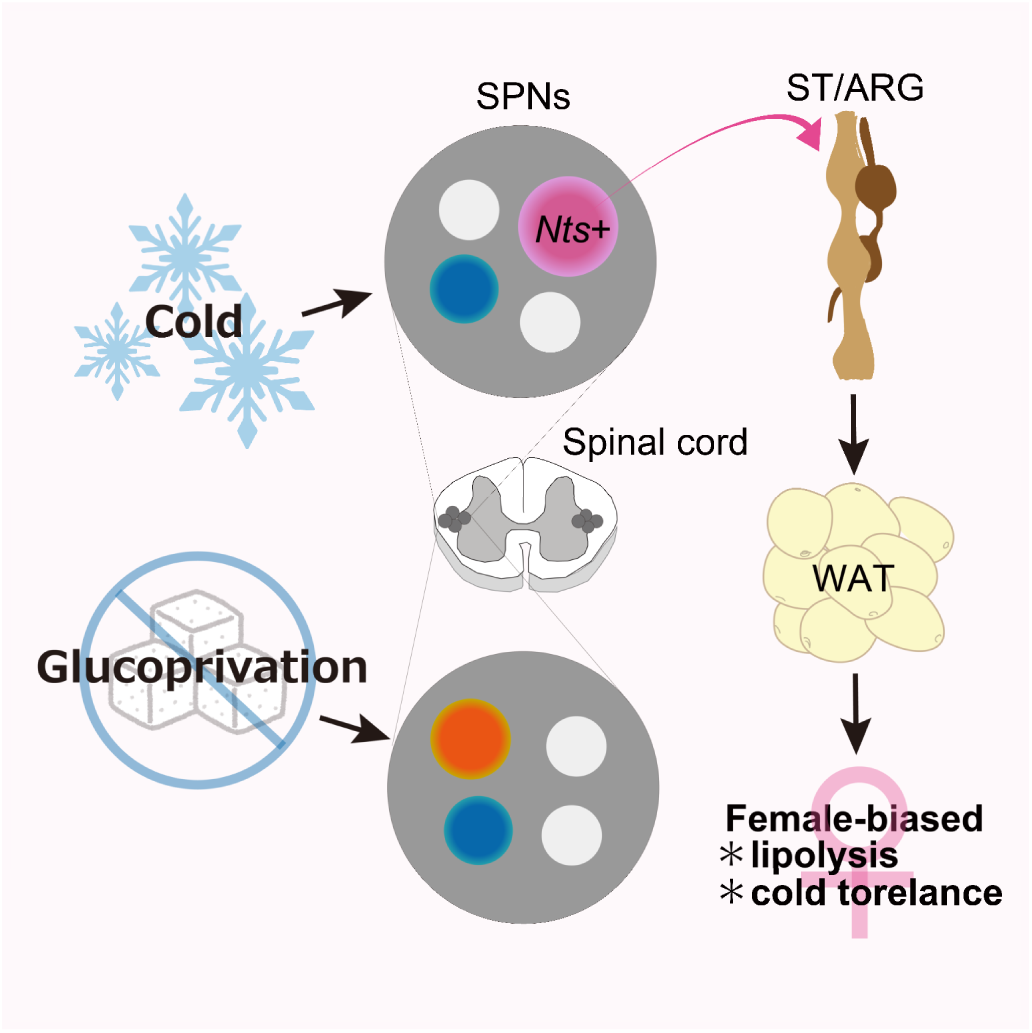

## Introduction

The sympathetic nervous system (SNS) coordinates the functions of multiple organs to maintain homeostasis and adapt to physiological stress^1–3^. It consists of two hierarchical layers: cholinergic spinal sympathetic preganglionic neurons (SPNs), which innervate postganglionic neurons in the sympathetic trunk and prevertebral ganglia, as well as the adrenal medulla^1^. Postganglionic neurons provide noradrenergic inputs to various peripheral organs. Although traditionally associated with a generalized “fight-or-flight” response^4^, sympathetic outflow can be highly selective across organs and physiological states^1,3^. Recent studies have begun to reveal a cellular basis for this selectivity in the peripheral SNS: molecularly defined postganglionic neuron types innervate distinct sets of organs or organ subdomains and regulate specific aspects of gastrointestinal, metabolic, and thermogenic physiology^5–9^. How such selective peripheral outputs are organized at the spinal preganglionic level, however, remains poorly understood.

SPNs occupy a pivotal position between descending autonomic commands and peripheral sympathetic ganglia. Recent single-nucleus RNA-sequencing studies have revealed more than a dozen molecularly distinct SPN subtypes^10,11^, and our previous work showed that genetically defined SPN populations can form parallel pathways to different peripheral targets^12^. These observations raise a broader question: do transcriptomically defined SPN subtypes constitute functional output modules that couple particular physiological demands to selective peripheral responses? This question remains unresolved even for major sympathetic functions. For example, the molecular identities and recruitment patterns of SPNs that drive adipose tissue lipolysis, a major sympathetic response to thermal and metabolic challenges^13^, remain unknown. Resolving this problem requires linking SPN molecular identity to spatial position, physiological recruitment, and peripheral output.

Spatial transcriptomic approaches provide an opportunity to resolve molecular identity and anatomical position within a common cellular framework^14–16^. Combined with immediate early gene (IEG) mapping, they can further link defined cell populations to activity associated with specific behavioral or physiological states^17,18^. They have been applied extensively in the brain and increasingly to sensory and motor populations of the spinal cord^19–23^, but the spatial and functional organization of spinal neurons controlling visceral organs remains largely unexplored. Such an integrated approach is particularly valuable for SPNs, which are sparse, dispersed, and heterogeneous^1,10–12^. It remains unknown whether physiological challenges broadly activate sympathetic output or instead recruit defined combinations of molecularly and spatially organized SPN populations.

Here, we generated a spatial transcriptomic activity atlas of mouse SPNs using the Xenium platform^24^ and examined their recruitment by two distinct homeostatic challenges: cold exposure, which drives sympathetic energy mobilization and thermogenesis^25,26^, and glucoprivation induced by 2-deoxyglucose (2-DG), which evokes sympathoadrenal responses that restore circulating glucose^12,27^. This framework allowed us to test whether physiological demands are represented by defined SPN output modules rather than by global sympathetic activation.

## Results

### Spatial organization of SPN transcriptomic subtypes

We designed 100 custom Xenium probes to cover representative marker genes for SPN subtypes, which were reclassified based on publicly available snRNA-seq data^10^ (Figure S1A, B), along with a panel of major IEGs. Using this custom probe set (Table S1) together with the predesigned Xenium Mouse Brain Gene Expression Panel (247 genes), we performed Xenium-based STx on 994 coronal spinal cord sections from 18 mice (nine males and nine females), spanning the thoracic, lumbar, and sacral segments (Figure 1A). After quality control and cell segmentation, approximately 3.2 million spinal cord cells were obtained. Through a series of classification steps, we identified 6,456 SPNs (0.2% of all detected cells) and classified them into 18 transcriptomic subtypes, each characterized by representative marker genes (Figures 1B–D and S1C, D). A representative coronal section (Figure 1C) confirmed that the detected SPNs were localized in the intermediolateral nucleus (IML), intercalated nucleus (IC), and central autonomic area (CA), consistent with their known distribution in the spinal cord^1^. We integrated the STx data with the snRNA-seq datasets using the robust cell-type decomposition (RCTD) method^28^ and found that most snRNA-seq-defined subtypes corresponded to a single STx-based subtype, supporting broad concordance between the two classifications (Figure S1E, F).

**Figure 1:**
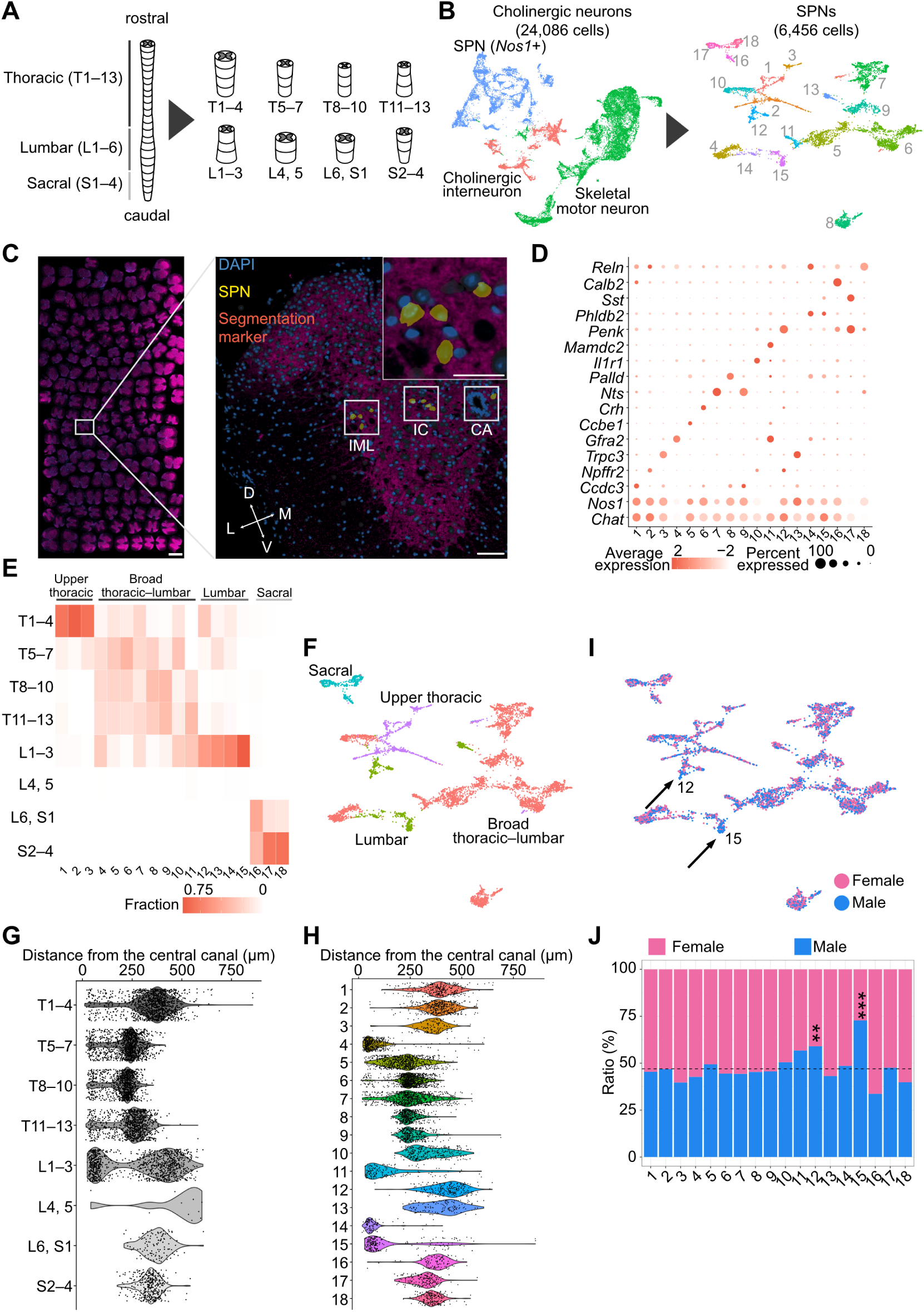
Spatial architecture of SPN transcriptomic subtypes. (A) Experimental scheme. (B) Uniform Manifold Approximation and Projection (UMAP) representations of cholinergic neurons (left) and SPNs (right). Dots are colored to indicate the three cholinergic populations (left) or 18 SPN clusters (right). (C) Fluorescent immunostaining images of a whole slide (left), a single spinal cord section (right), and a magnified view of the IML (inset), stained for segmentation markers (ATP1A1, E-cadherin, and CD45; magenta) with DAPI (blue). Yellow regions indicate SPNs. D, dorsal; V, ventral; L, lateral; M, medial. Scale bars: 1 mm (left), 100 μm (right), and 50 μm (inset). (D) Dot plot showing representative marker genes for each SPN subtype. (E) Heatmap showing the relative distribution of each SPN subtype across spinal segments. Four rostrocaudal distribution patterns are indicated: upper thoracic-specific (clusters 1–3), broad thoracic to lumbar (clusters 4–11), lumbar-specific (clusters 12–15), and sacral-specific (clusters 16–18). (F) UMAP of SPNs colored by the four rostrocaudal distribution patterns. (G, H) Violin plots showing the distance between SPNs and the central canal by spinal segment (G) and by SPN subtype (H). (I) UMAP of SPNs colored by sex. (J) Bar graph showing the proportion of each SPN subtype by sex. The dashed line indicates the overall male-to-female ratio of all SPNs. \*\**p* < 0.01, \*\*\**p* < 0.001, by Fisher’s exact test with Holm’s correction. For more data, see Figure S1 and Source Data file.

Next, we examined the distribution of SPN subtypes along the rostrocaudal axis, because rostrocaudal SPN position broadly reflects the organotopic organization of sympathetic outflow^1^. We noted that SPNs were sparse in the 4^th^ lumbar (L4)–L5 segments. Unbiased *k*-means clustering revealed four distributional patterns: the upper thoracic (T)-enriched group (three subtypes), lumbar-enriched group (four subtypes), sacral-specific group (three subtypes), and thoracic-to-lumbar distributed group (eight subtypes) (Figures 1E and S1G). These broad patterns were well separated in transcriptomic space (Figure 1F), indicating that distinct SPN transcriptomic subtypes occupy characteristic rostrocaudal positions within the spinal cord.

We also investigated the organization of SPNs along the mediolateral axis of the spinal cord by measuring the distance between SPNs and the central canal (Figure 1G, H). SPNs were broadly distributed but showed regional bias, being primarily localized to the IML in the thoracic segments, bimodally distributed between the IC and IML in the upper lumbar segments, and almost exclusively restricted to the IML in the sacral segments. Notably, several SPN subtypes were exclusively located in the CA, whereas others were predominantly located in the IML (Figure 1H). These findings demonstrate that mediolateral positioning within the spinal cord is also represented in the transcriptomic organization of SPNs.

Sexual dimorphism in SPNs has been described in rats, most notably in the lumbar segments, where SPNs project through the hypogastric nerve to regulate the pelvic autonomic ganglia and urogenital organs^29,30^. Therefore, we investigated whether our dataset contained sex-specific or -biased SPN subtypes. Although all SPN subtypes were present in both sexes, we found a significant male bias in the abundance of cluster 12, marked by *proenkephalin* (*Penk*), and cluster 15, marked by *somatostatin* (*Sst*) (Figure 1I, J), which were almost exclusively distributed in the upper lumbar segments (Figure 1E). Thus, the transcriptomic organization of SPNs also exhibits sex-biased features, most prominently among the upper-lumbar subtypes.

### Distinct stressors recruit different combinations of SPN transcriptomic subtypes

Because multiple SPN transcriptomic subtypes are intermingled within common spinal segments (Figure 1E), it is important to determine whether a given stressor recruits the entire local SPN population (Figure 2A_1_) or selectively engages defined subtypes, either in accordance with their transcriptomic identity (Figure 2A_2_) or independently of it (Figure 2A_3_). This distinction is essential to understand how central commands are conveyed to peripheral targets through transcriptomically defined SPN subtypes. To distinguish among these possibilities, we examined the IEG expression in SPNs under cold exposure and glucoprivation conditions (Figure 2B). Our dataset also included mice maintained at room temperature (RT) and injected with saline as controls. Although *c-Fos* expression was minimal in the control group, both 1.5 h cold exposure and 2-DG administration induced *c-Fos* expression in subsets of SPNs (Figure 2C), arguing against uniform activation. The cold-responsive population was enriched in two SPN transcriptomic subtypes characterized by *corticotropin-releasing hormone* expression (*Crh*+; cluster 6) and *neurotensin* expression with low *nitric oxide synthase 1* expression (*Nts*+ *Nos1*^low^; cluster 7) (Figure 2D). In contrast, the glucoprivation-responsive population was dominated by *Crh*+ SPNs (cluster 6) and *Palladin*-expressing SPNs (*Palld*+; cluster 8). This correlation between *c-Fos* expression and SPN transcriptomic subtype identity was further supported by quantitative analysis (Figure 2E). These data indicate that physiological challenges recruit defined SPN transcriptomic subtypes, most closely matching model 2 in Figure 2A, with different stressors engaging partially overlapping yet distinct combinations of SPN transcriptomic subtypes.

**Figure 2:**
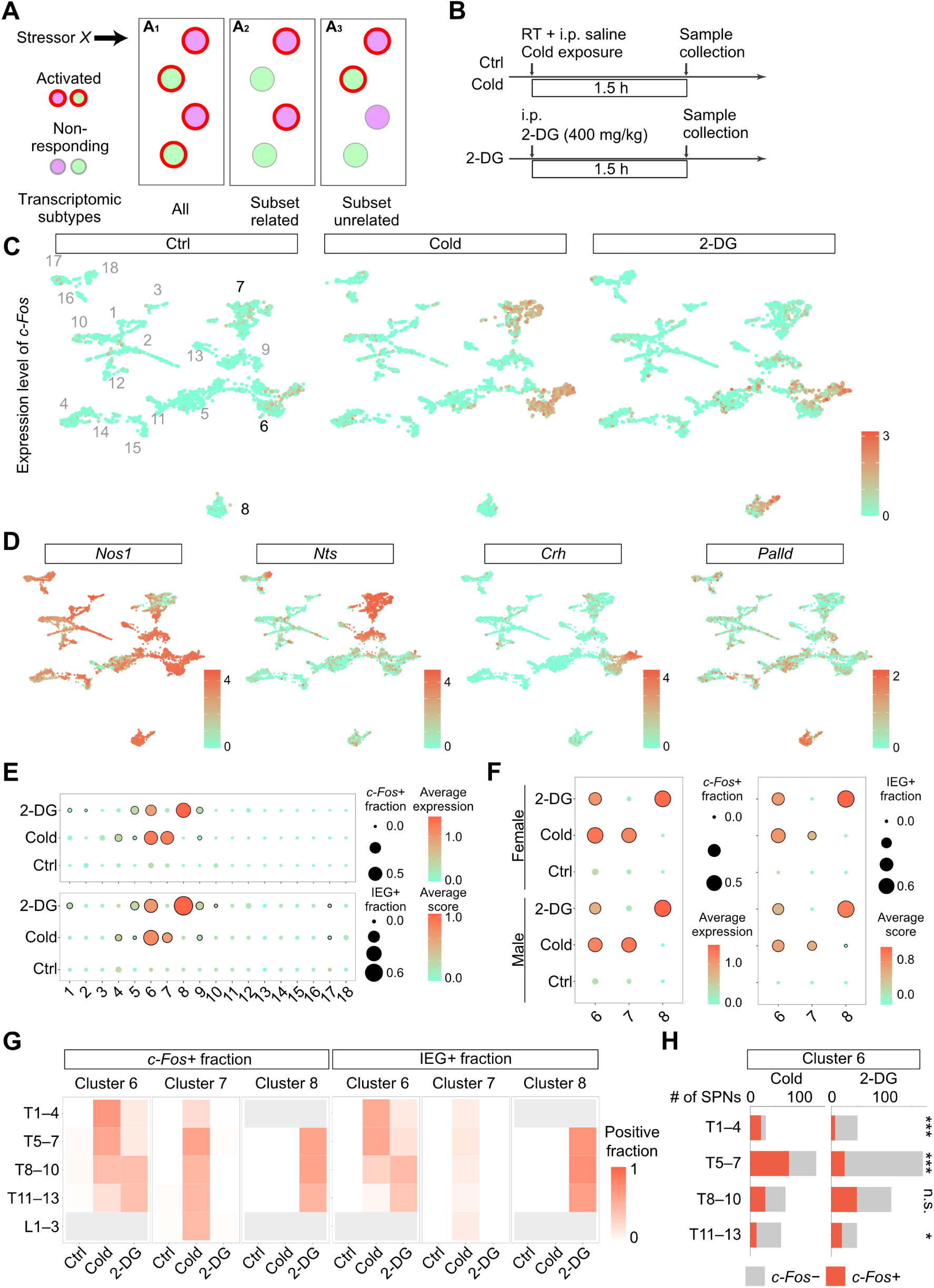
Physiological stressors recruit distinct combinations of SPN transcriptomic subtypes. (A) Three possible relationships between stressor *X*-induced SPN activation and transcriptomic subtype identity. Purple and green circles indicate distinct SPN transcriptomic subtypes, and red contours denote activated cells. In A_1_, all local SPNs are activated. In A_2_, activation is restricted to a transcriptomically defined subset, here the purple population. In A_3_, activation occurs in a subset of SPNs independently of transcriptomic subtype identity. (B) Experimental scheme. RT, room temperature; i.p., intraperitoneal injection; Ctrl, control group. (C, D) UMAP plots showing *c-Fos* expression under each condition (C) and expression of the indicated marker genes (D) across all SPNs. (E) Dot plots showing *c-Fos* expression level and *c-Fos*+ fraction (top), and IEG score and IEG+ fraction (bottom), across SPN subtypes. (F) Dot plots showing *c-Fos* expression level and *c-Fos*+ fraction (left), and IEG score and IEG+ fraction (right), for clusters 6–8 in each sex. In (E) and (F), dots outlined in black indicate expression levels significantly higher than those in the RT-saline control group, as determined by the Wilcoxon rank-sum test with Holm’s correction (*p* < 0.05). (G) Heatmaps showing the *c-Fos*+ (left) or IEG+ (right) fraction across thoracic–lumbar segments for clusters 6–8 under each condition. For each subtype, segments containing fewer than 5% of all SPNs within the cluster were excluded from the display (shown in gray). (H) Segmental distribution of cluster 6 (*Crh*+ SPNs) and the *c-Fos*+ fraction following cold exposure or 2-DG administration. \*\*\**p* < 0.001, \**p* < 0.05, and n.s. (not significant), determined by Fisher’s exact test with Holm’s correction. For more data, see Figure S2 and Source Data file.

Cold exposure and glucoprivation induced not only *c-Fos* but also multiple IEGs, including *Fosb*, *Nr4a1*, and *Npas4* (Figure S2A). Therefore, we considered multiple IEGs together to define an IEG score (see Methods) that supported the cold- and glucoprivation-responsive SPNs identified using *c-Fos* (Figure S2B). In addition, because integrating multiple IEGs improved the signal-to-noise ratio, the IEG score revealed an additional weak activation of SPN transcriptomic subtypes (Figure 2E, bottom). Within clusters 6, 7, and 8, the induction of multiple IEGs was observed in response to the corresponding stressors (Figure S2C). No apparent sex differences were observed in activation patterns (Figures 2F and S2D). Collectively, these findings further support the stressor-selective recruitment of specific SPN subtypes in both sexes.

Because clusters 6–8 belonged to the broad thoracic-to-lumbar distribution group (Figure 1E), we next investigated whether IEG induction occurred in specific spinal segments. Within cluster 6 (*Crh*+ SPNs), cold exposure tended to activate SPNs in the upper thoracic segments (T1–T7), whereas glucoprivation predominantly induced IEGs at lower thoracic levels (T8–T13) (Figure 2G, H). In contrast, in clusters 7 and 8, the stressor-responsive SPNs were largely distributed in proportion to their overall rostrocaudal distribution (Figure 2G) in both sexes (Figure S2E). These data suggest that SPN recruitment reflects both transcriptomic subtype identity and spinal location.

Taken together, these data show that physiological challenges recruit distinct combinations of molecularly and spatially organized SPN populations, supporting a modular organization of sympathetic control at the spinal preganglionic level.

### Molecularly defined SPNs preferentially engage distinct sympathetic output pathways

If these stressor-selective SPN populations constitute functional modules, they should differ not only in when they are recruited, but also in where their activity is routed peripherally. We tested this prediction by focusing on lower-thoracic *Nts*+ SPNs, which included the cold-responsive cluster 7. Using histochemistry, we examined their identity and cold responsiveness in the T8–T12 segments (Figure 3A). *Nts*+ SPNs were predominantly located in the IML, sparsely distributed in the IC, and absent from the CA region (Figure S3A), consistent with Figure 1H. Within the IML, *Nts*+ SPNs accounted for approximately one-third of the choline acetyltransferase (ChAT)+ pan-SPN population, with no detectable sex difference, consistent with Figure 1J. Following cold exposure, *c-Fos* was induced in approximately 40% of SPNs in the IML (Figure S3B), and the majority of *Nts*+ SPNs became *c-Fos*+, representing ∼50% of all *c-Fos*+ SPNs in this region (Figures 3B, C, and S3C). These results independently validate the STx-based activity atlas and identify lower-thoracic *Nts*+ SPNs as a major cold-responsive SPN population.

**Figure 3:**
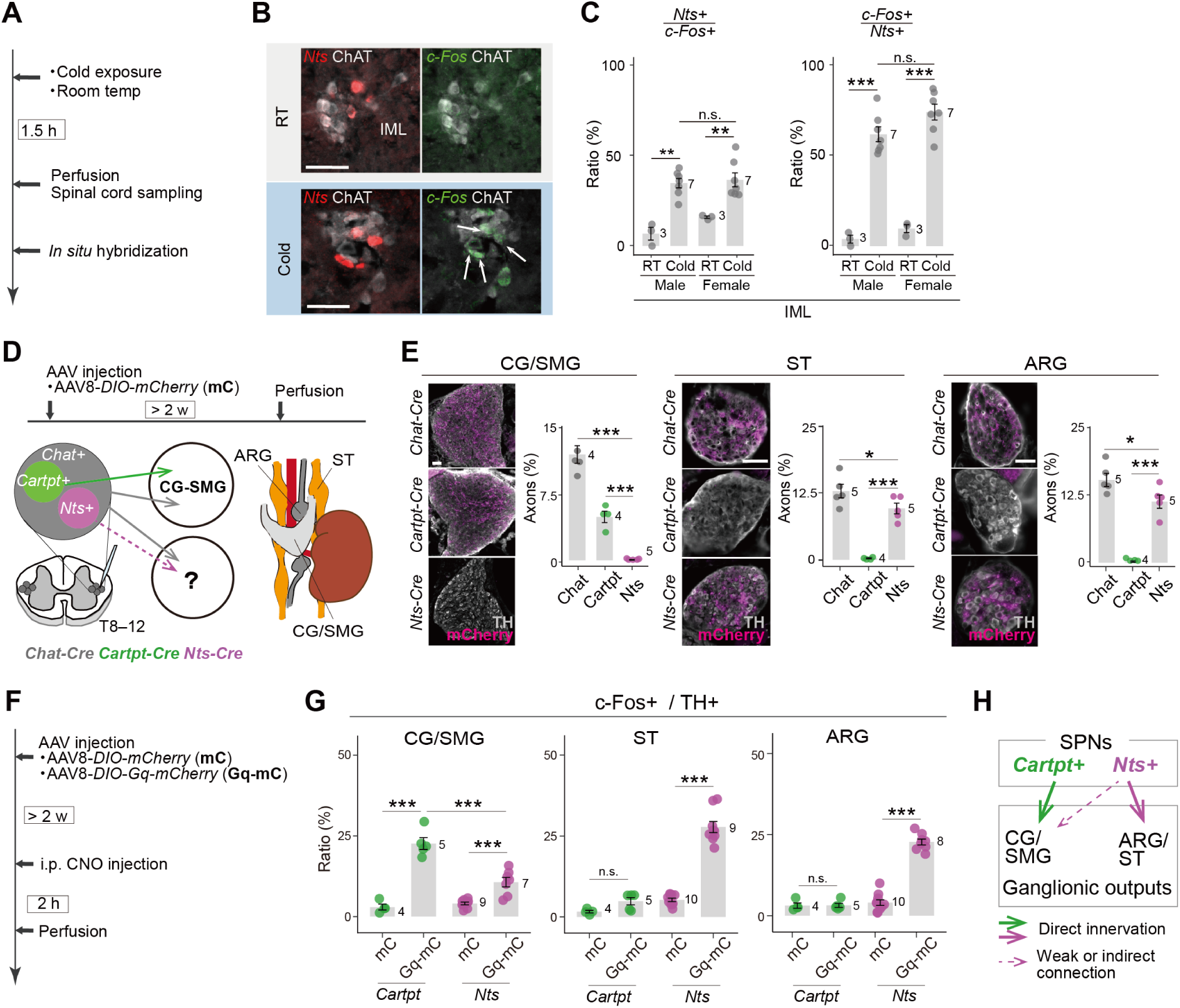
Molecularly defined SPNs preferentially engage distinct sympathetic ganglionic outputs. (A) Experimental scheme and timeline for cold exposure. (B) Representative images of the IML showing *Nts* (red), *c-Fos* (green), and ChAT (gray) in the RT (top) and cold (bottom) exposure groups. Arrows indicate *c-Fos*+ *Nts*+ ChAT+ triple-positive cells. (C) Quantification of the fraction of *Nts+* cells among *c-Fos+* cells (left) and *c-Fos+* cells among *Nts+* cells (right) in the IML of the RT and cold exposure groups in both sexes. One-way ANOVA followed by Tukey’s post hoc test: \*\**p* < 0.01, \*\*\**p* < 0.001. (D) Schematic of the experimental design and timeline. (E) Proportion of mCherry+ axonal area within TH+ regions originating from *Chat*+, *Cartpt*+, and *Nts*+ SPNs in the CG/SMG, ST, and ARG. Representative sections (left) and quantification (right). One-way ANOVA with post hoc Tukey’s test: \**p* < 0.05, \*\*\**p* < 0.001. (F) Schematic of the timeline for chemogenetic activation of the SPNs and c-Fos analysis in the postganglionic neurons. (G) Fraction of c-Fos+ cells among TH+ cells in the CG/SMG, ST, and ARG following chemogenetic activation of *Cartpt*+ or *Nts*+ SPNs. One-way ANOVA with post hoc Tukey’s test: n.s., not significant; \*\*\**p* < 0.001. (H) Schematic summary of the ganglionic outputs of *Cartpt*+ and *Nts*+ SPNs. The number of animals in each group is shown. Both sexes were included in the analyses. Error bars represent the standard deviation. Scale bars, 50 μm. For more data, see Figures S3 and S4.

We next asked whether this cold-responsive population engages a characteristic sympathetic output pathway. To analyze the axonal projection targets of *Nts*+ SPNs, we injected an AAV expressing Cre-dependent mCherry into the T8–T12 spinal segments of *Nts-Cre* mice^31^, using *Chat-Cre*^32^ (a pan-SPN driver) and *Cartpt-Cre* (a CG/SMG-projecting SPN marker^12^) mice as references (Figure 3D). Dual-color histochemical analysis in wild-type mice showed that *Nts*+ and *Cartpt*+ SPNs were largely non-overlapping (Figure S4A), consistent with our STx dataset showing enrichment of *Cartpt* expression in transcriptomic clusters distinct from the major *Nts*+ population (Figure S1H). Histochemical analyses following viral injection confirmed efficient and selective viral targeting of *Nts*+ SPNs within the IML, with mCherry expressed in the majority of *Nts*+ SPNs and little labeling of Nts− SPNs (Figure S4B). We also detected *Nts* expression and mCherry labeling outside SPNs, particularly in putative sensory neurons in the dorsal horn (Figure S4C, D; see Supplementary Note 1).

Two to three weeks after viral injection, *Chat*+ SPN axons were detected in the CG/SMG and lower thoracic sympathetic trunk (ST). Consistent with our previous findings^12^, *Cartpt+* SPNs preferentially targeted the CG/SMG and showed little projection to the ST. In contrast, *Nts*+ SPNs showed minimal axonal labeling in the CG/SMG but dense projections to the ST at T8–T12 (Figures 3E, middle and S3D). In addition, *Nts*+ SPNs sent prominent axonal projections to the aorticorenal ganglia (ARG), which received little *Cartpt*+ SPN input (Figure 3E, right). Neither population showed substantial innervation of the adrenal medulla (Figure S3E), which we previously identified as a major target of *Oxtr*+ (*Palld*+) SPNs^12^. The axonal projections of *Nts*+ SPNs to the ST and ARG did not differ significantly between male and female mice (Figure S3F). Thus, *Nts*+ and *Cartpt*+ SPNs exhibit strongly biased and distinct patterns of sympathetic ganglionic innervation.

We next asked whether these anatomical differences correspond to distinct patterns of postganglionic activation. We targeted the chemogenetic activator hM3Dq-mCherry to *Cartpt*+ or *Nts*+ SPNs in the T8–T12 segments (Figure 3F). More than two weeks later, clozapine-N-oxide (CNO)-mediated activation of these SPNs induced c-Fos in tyrosine hydroxylase (TH)+ postganglionic neurons. Activation of *Cartpt*+ SPNs significantly increased c-Fos in the CG/SMG but not in the ST or ARG (Figure 3G). In contrast, activation of *Nts*+ SPNs produced only a weak response in the CG/SMG but robustly induced c-Fos in the ST and ARG, with no detectable sex difference (Figure S3G). The weak CG/SMG response may reflect sparse direct projections below our detection threshold or indirect activation through downstream physiological crosstalk. Collectively, these data functionally show that molecularly defined SPN populations preferentially recruit distinct and biased patterns of sympathetic ganglionic activation (Figure 3H).

Having established that *Nts*+ SPNs preferentially engage the ST and ARG, we next asked which organ-projecting postganglionic populations are recruited through this pathway. Because the ARG contains postganglionic neurons innervating the kidney^33,34^ and WAT^7,35,36^, we first examined their organization using dual-color retrograde labeling with cholera toxin subunit B (CTb) from combinations of the kidney, subcutaneous inguinal WAT (iWAT), and gonadal WAT (gWAT) (Figure 4A). When the two tracers were mixed and injected into iWAT, the vast majority of labeled postganglionic neurons in the ST and ARG were co-labeled, validating the dual-tracing approach. In contrast, labeling from distinct organs showed largely separate kidney- and WAT-projecting populations, with only limited overlap between iWAT- and gWAT-projecting neurons (Figure 4B).

**Figure 4:**
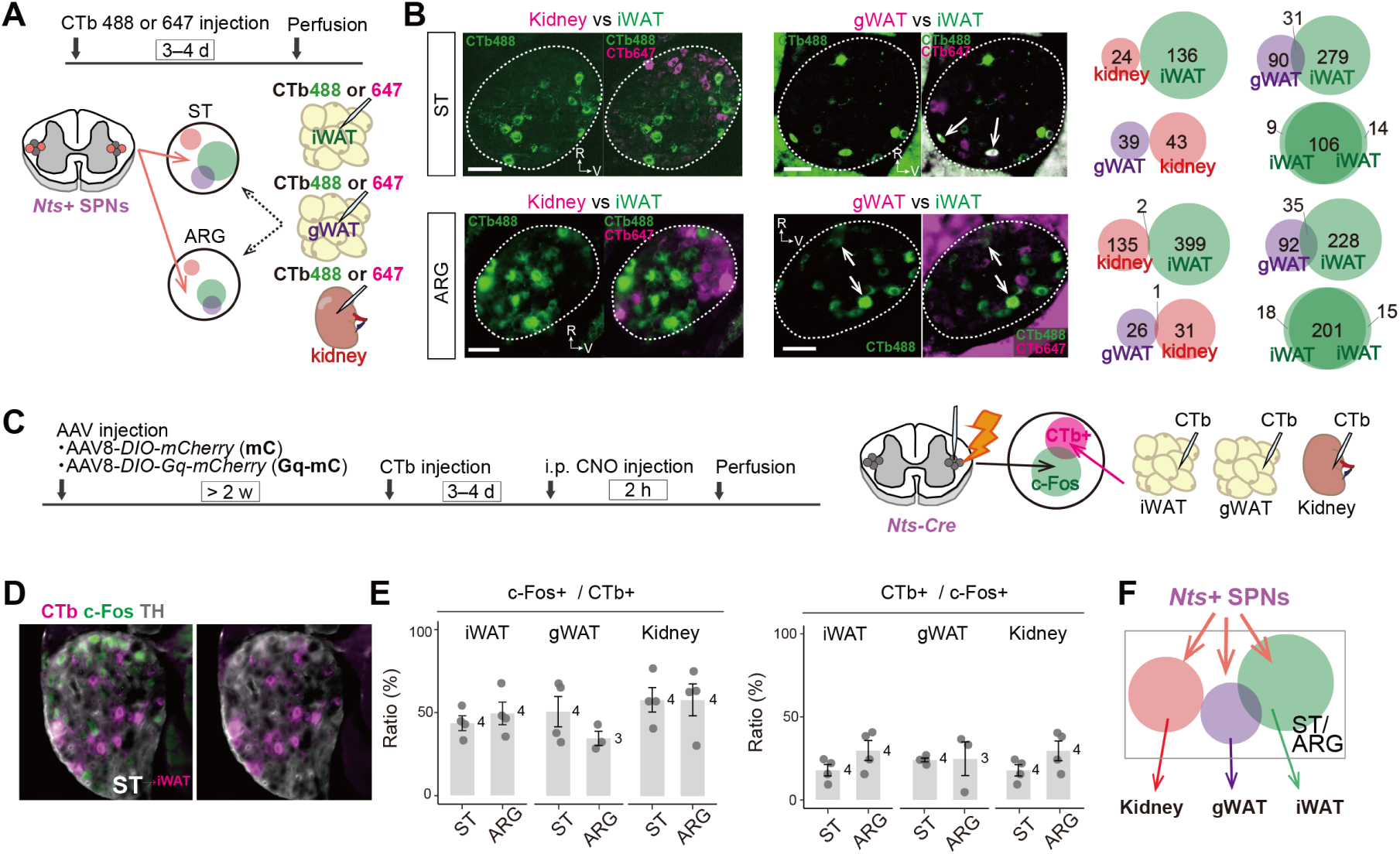
*Nts*+ SPNs recruit postganglionic pathways to WAT and kidney. (A) Schematic of the experimental design for dual-color retrograde tracing. (B) Left, representative images of the ST (top) and ARG (bottom) showing postganglionic neurons retrogradely labeled from the indicated organs. R, rostral; V, ventral. Arrows indicate dual-positive cells. Right, quantification of CTb-labeled postganglionic neurons in the ST (top) and ARG (bottom). Data from N = 3–7 mice were pooled. (C) Experimental timeline for chemogenetic activation of *Nts*+ SPNs to examine c-Fos induction in postganglionic neurons innervating iWAT, gWAT, and kidney. Some animals used in this experiment also contributed to the analyses shown in Figure 3G. (D) Representative images of the ST showing c-Fos (green), TH (gray), and CTb-labeled postganglionic neurons (magenta) retrogradely traced from iWAT. (E) Fraction of c-Fos+ cells among CTb+ cells (left) and CTb+ cells among c-Fos+ cells (right) in the ST and ARG following chemogenetic activation of *Nts*+ SPNs. (F) Schematic summary of organ-projecting postganglionic populations recruited downstream of *Nts*+ SPNs. The number of animals in each group is shown. Both sexes were included in the analyses. Error bars represent the standard deviation. Scale bars, 50 μm.

We next asked whether these organ-projecting populations are recruited by *Nts*+ SPNs. We targeted hM3Dq-mCherry to *Nts*+ SPNs in the T8–T12 segments and injected CTb into iWAT, gWAT, or the kidney (Figure 4C). CNO administration induced c-Fos in approximately half of the CTb-labeled postganglionic neurons in both the ST and ARG, irrespective of their peripheral target (Figure 4D and 4E, left). Conversely, CTb-labeled neurons accounted for ∼20% of all *c-Fos*+ postganglionic neurons for each individual target examined (Figure 4E, right). Thus, *Nts*+ SPNs recruit postganglionic pathways projecting to iWAT, gWAT, and kidney through the ST and ARG (Figure 4F), while leaving open the possibility of additional downstream targets.

### *Nts*+ SPNs regulate WAT metabolism in a female-biased manner

Given the recruitment of *Nts*+ SPNs during cold exposure and their engagement of postganglionic pathways to WAT, we hypothesized that *Nts*+ SPNs facilitate the mobilization of stored lipids to support adaptation to cold. We first tested this prediction by chemogenetically activating *Nts*+ SPNs and measuring plasma non-esterified fatty acid (NEFA) levels (Figure 5A).

**Figure 5:**
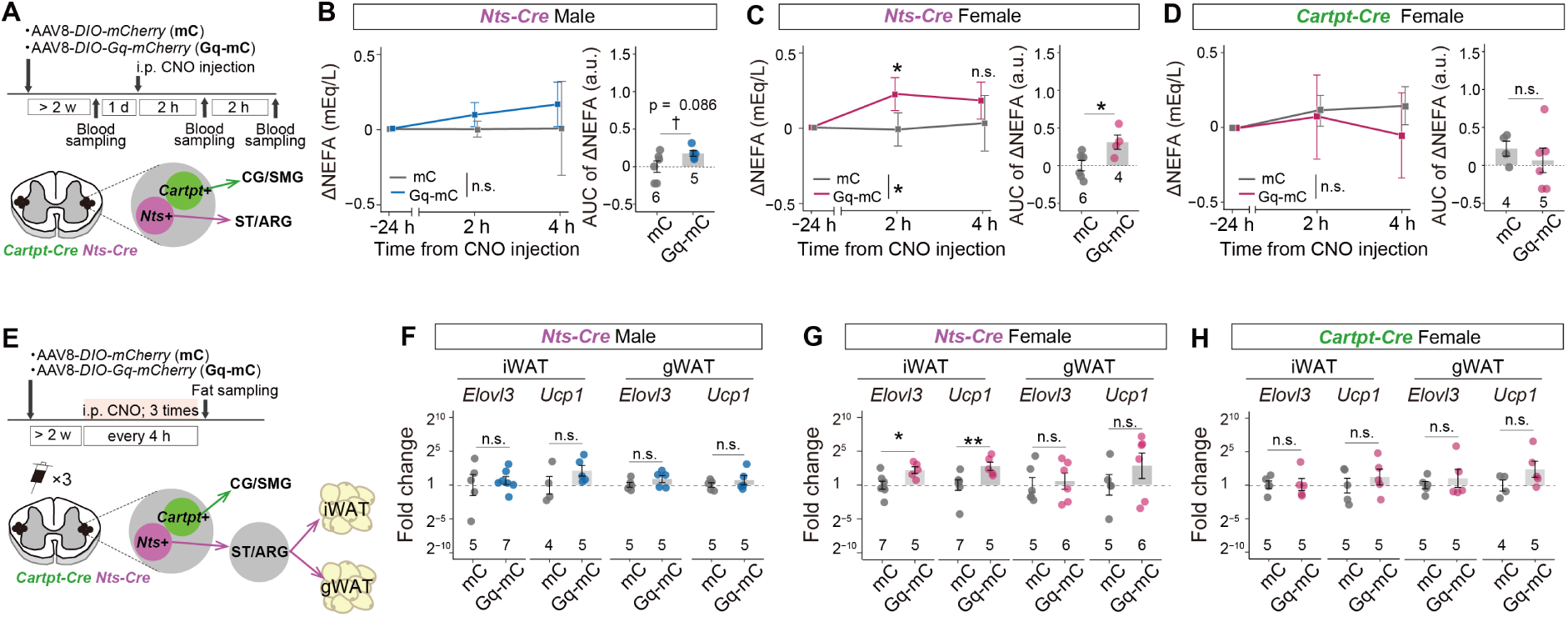
*Nts+* SPN activation promotes female-biased lipid mobilization. (A) Experimental scheme for plasma NEFA measurements following chemogenetic activation of *Nts*+ or *Cartpt*+ SPNs. (B–D) Changes in plasma NEFA levels after CNO injection relative to the previous day in *Nts-Cre* males (B), *Nts-Cre* females (C), and *Cartpt-Cre* females (D). Right panels show the AUC of ΔNEFA over 4 h after CNO injection in control (mCherry, mC) and Gq-injected (Gq-mC) mice. Two-way repeated-measures ANOVA: group, time, and interaction effects, n.s. (B, D); group effect, \**p* < 0.05, and time and interaction effects, n.s. (C). Group differences at each time point were analyzed post hoc using a two-sided unpaired t-test with Bonferroni correction (\**p* < 0.05). AUC data were analyzed using a two-sided Welch’s t-test (n.s., not significant; †*p* < 0.1; \**p* < 0.05). (E) Experimental scheme for quantitative PCR analysis following three repeated chemogenetic activations of *Nts*+ or *Cartpt*+ SPNs. (F–H) Fold changes in *Elovl3* and *Ucp1* expression in iWAT and gWAT from control (mC) and Gq-mC *Nts-Cre* males (F), *Nts-Cre* females (G), and *Cartpt-Cre* females (H). \**p <* 0.05*;**p <* 0.01 by the Wilcoxon rank-sum test. n.s., not significant. The number of animals in each group is shown. Error bars represent the standard deviation. Blue and pink dots represent male and female data, respectively. For more data, see Figure S4.

In males, NEFA levels tended to increase following *Nts*+ SPN activation, but the effect did not reach statistical significance (Figure 5B). In contrast, activation of *Nts*+ SPNs in females significantly increased plasma NEFA levels, whereas control mice expressing mCherry alone showed no change (Figure 5C). To determine whether this response was specific to the *Nts*+ SPN pathway, we similarly activated *Cartpt*+ SPNs in females (Figure S4E), which did not alter NEFA levels (Figures 5D). Thus, *Nts*+ SPNs exert a female-biased effect on lipid mobilization that is not reproduced by activation of *Cartpt*+ SPNs, consistent with their distinct ganglionic output patterns (Figure 3E, G).

We next examined adipose transcriptional responses following repeated SPN activation, focusing on *Elovl3* and *uncoupling protein 1* (*Ucp1*), genes associated with sympathetic and thermogenic activation of WAT^37,38^ (Figure 5E). Three consecutive activations of *Nts*+ SPNs within a single day did not significantly alter *Elovl3* or *Ucp1* expression in iWAT or gWAT of males (Figure 5F). In females, however, the same manipulation significantly increased *Elovl3* and *Ucp1* expression in iWAT, with less pronounced changes in gWAT (Figure 5G). This preferential effect on iWAT may reflect the greater representation of iWAT-projecting postganglionic neurons within the ST and ARG examined here (Figure 4B). Activation of *Cartpt*+ SPNs in females did not significantly alter *Elovl3* or *Ucp1* expression in either iWAT or gWAT (Figure 5H). Together, these data show that *Nts*+ SPNs promote female-biased lipid mobilization accompanied by adaptive transcriptional responses in iWAT.

To determine whether sustained *Nts*+ SPN activation produces longer-term changes in adipose stores, we administered CNO once daily for seven consecutive days (Figure 6A). Overall body weight remained unchanged in both sexes, although males showed a modest increase in food intake, whereas females did not (Figure 6B, C). Repeated activation of *Nts*+ SPNs markedly reduced the relative mass of both iWAT and gWAT in females but not in males (Figure 6D, E, I). Consistent with this reduction, adipocytes in both depots were significantly smaller in females, with their size distributions shifted toward smaller cells (Figure 6F–H, 6J–L). In males, the effect was limited to a modest reduction in iWAT adipocyte size. Thus, repeated *Nts*+ SPN activation depletes WAT stores in a strongly female-biased manner.

**Figure 6:**
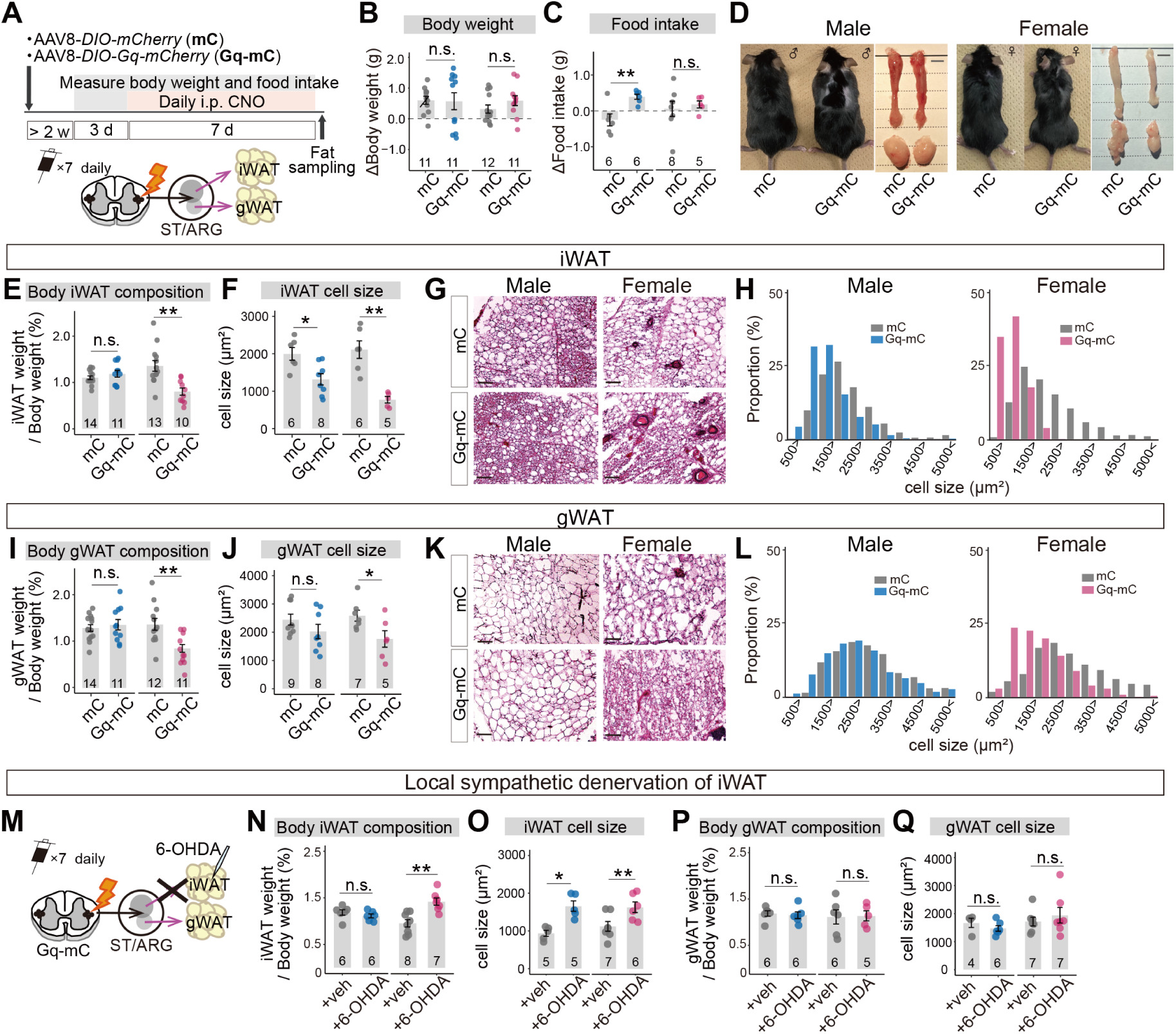
Repeated *Nts*+ SPN activation remodels WAT via local sympathetic innervation. (A) Experimental scheme for chronic daily chemogenetic activation of *Nts*+ SPNs. (B, C) Changes in body weight (B) and cumulative food intake (C) over the 7-day activation period. (D) Representative images of male (left) and female (right) mice and dissected iWAT and gWAT after chronic activation. Scale bar, 5 mm. (E, I) Relative mass of iWAT (E) and gWAT (I). (F, J) Quantification of adipocyte size in iWAT (F) and gWAT (J). (G, K) Representative H&E-stained images of iWAT (G) and gWAT (K) from control (top) and Gq-mC (bottom) mice of both sexes. Scale bars, 50 μm. (H, L) Distribution of adipocyte size in iWAT (H) and gWAT (L).(M) Schematic of the experimental scheme for local sympathetic denervation of iWAT during chronic chemogenetic activation of *Nts*+ SPNs. (N–Q) Relative mass of iWAT (N) and gWAT (P), and quantification of adipocyte size in iWAT (O) and gWAT (Q), in vehicle-treated (+veh) and 6-OHDA-treated (+6-OHDA) groups. \**p* < 0.05, \*\**p* < 0.01 by two-sided Welch’s t-test. The number of animals in each group is shown. Error bars represent the standard deviation. Blue and pink dots represent male and female data, respectively.

We next asked whether this adipose phenotype requires intact local sympathetic innervation. In an independent cohort, we chemically denervated iWAT with 6-hydroxydopamine (6-OHDA) during repeated chemogenetic activation of *Nts*+ SPNs of female mice (Figure 6M). Local denervation prevented the reduction in iWAT mass and adipocyte size observed after repeated *Nts*+ SPN activation (Figure 6N, O). By contrast, 6-OHDA did not alter gWAT mass or adipocyte size (Figure 6P, Q), consistent with a local effect of iWAT denervation. These results indicate that the chronic iWAT phenotype induced by *Nts*+ SPN activation depends on intact local sympathetic innervation. Together with the acute lipid mobilization observed above, these data establish a female-biased pathway through which *Nts*+ SPNs promote the mobilization and depletion of WAT lipid stores.

### *Nts*+ SPNs confer cold tolerance in female mice

To determine whether *Nts*+ SPNs are required for physiological adaptation to cold, we next examined their loss-of-function effects. We bilaterally injected an AAV expressing Cre-dependent active caspase (taCasp3-TEV)^39^ or mCherry (control) into the T8–T12 spinal cord segments of *Nts-Cre* mice (Figure 7A). After recovery, mice were subjected to mild fasting for 6 h followed by 4°C cold exposure for 4 h. Under these conditions, cold exposure robustly induced *c-Fos* in *Nts*+ SPNs in both sexes (Figure S5A, B). Post hoc histochemical analyses confirmed a significant reduction in *Nts*+ SPNs in the IML of the lower thoracic spinal cord in taCasp-expressing mice (Figure 7B, C), without affecting *Nts*+ sensory neurons in the dorsal horn (Figure S5C).

**Figure 7:**
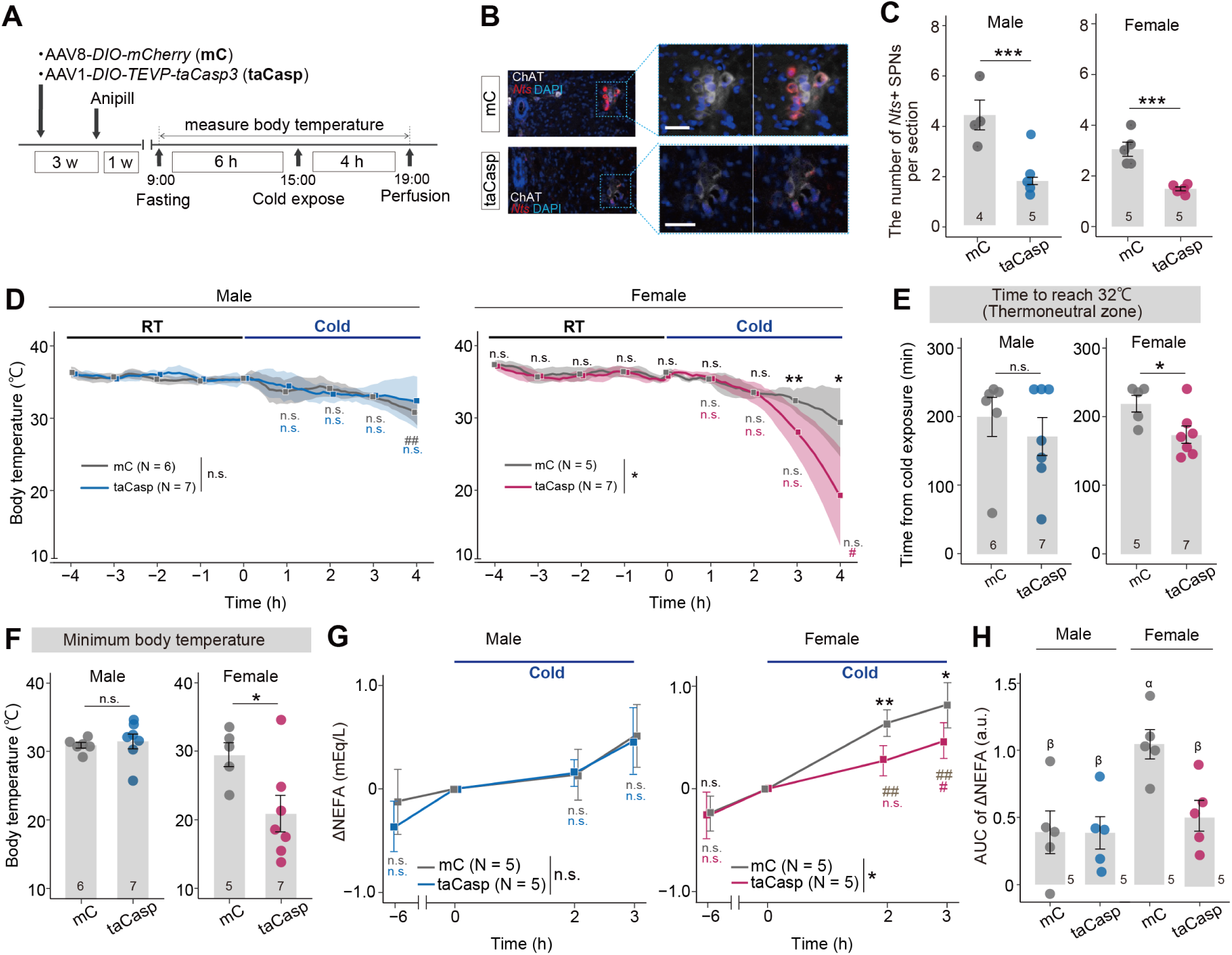
*Nts*+ SPNs confer cold tolerance in female mice without food access. (A) Experimental scheme for measuring Tc during cold exposure without food access. (B) Coronal spinal cord sections showing *Nts*+ SPNs (red), ChAT (gray), and DAPI (blue) in the IML of the control (mC; top) and taCasp (bottom) mice. Right panels show magnified, channel-separated views of the boxed regions. Scale bar, 50 μm. (C) Quantification of *Nts*+ SPN numbers per section in control and taCasp mice. *\*\*\*p <* 0.001 by two-sided Welch’s t-test. (D) Tc before and during cold exposure in male (left) and female (right) mice. Two-way repeated-measures ANOVA: group effect, n.s., time effect, *p* < 0.001, and interaction effect, n.s. (male); group effect, \**p* < 0.05, time effect, *p* < 0.001, and interaction effect, *p* < 0.05 (female). (E) Latency to reach a Tc of 32 °C during cold exposure. (F) Minimum Tc reached during the 4-h cold exposure. *\*p <* 0.05 by two-sided Welch’s t-test. (G) Changes in plasma NEFA levels (mEq/L) relative to the onset of cold exposure (time zero). Two-way repeated-measures ANOVA: group effect, n.s.; time effect, *p* < 0.001; and interaction effect, n.s. (male); group effect, \**p* < 0.05; time effect, *p* < 0.001; and interaction effect, \*\**p* < 0.01 (female). (H) AUC of ΔNEFA over 3 h of cold exposure. Different letters (α and β) indicate *p <* 0.05 by one-way ANOVA with Tukey’s post hoc test. In panels (D) and (G), the time effect relative to time 0 was analyzed using one-way ANOVA followed by Bonferroni’s multiple-comparisons test (#*p* < 0.05, ##*p* < 0.01; colors indicate groups). For the female data, group differences at each time point were further assessed post hoc using a two-sided unpaired t-test with Bonferroni correction (\**p* < 0.05, \*\**p* < 0.01, indicated at the top of the graph). The body temperature measurements in (D–F) and plasma NEFA measurements in (G, H) were performed in separate cohorts of mice. The number of animals in each group is shown. Error bars and shaded areas (D) represent the standard deviation. For more data, see Figures S5–S7.

We then monitored core body temperature (Tc) continuously during the cold challenge. Control mice of both sexes maintained Tc relatively well throughout the 4-h cold exposure (Figure 7D). Ablation of *Nts*+ SPNs had little effect in males, whereas females exhibited a marked decline in Tc, reached 32°C more rapidly, and attained a lower minimum Tc than control females (Figure 7D–F). In several ablated females, Tc fell rapidly to below 20°C, demonstrating a severe failure to maintain body temperature during cold exposure. These data suggest a substantial impairment of cold tolerance. Importantly, c-Fos induction in central cold-responsive and thermogenic neurons^40–42^ remained intact after *Nts*+ SPN ablation (Figure S5D, E), arguing against a major defect in upstream central cold sensing and supporting a deficit at the sympathetic efferent level.

Because activation of *Nts*+ SPNs promoted lipid mobilization in females (Figures 5 and 6), we next asked whether impaired cold tolerance was accompanied by defective mobilization of stored lipids. Cold exposure increased plasma NEFA levels in control mice, with a larger response in females than in males (Figure 7G, H). Ablation of *Nts*+ SPNs markedly attenuated this cold-induced NEFA response in females but had little effect in males. Thus, the sex-biased thermoregulatory phenotype was paralleled by a selective impairment of lipid mobilization in females.

To assess whether chronic ablation itself altered adipose tissue, we examined additional cohort of female mice eight weeks after viral injection. In the absence of cold exposure, *Nts*+ SPN ablation produced no overt changes in iWAT or gWAT mass, adipocyte size, or expression of adipose metabolic genes (Figure S6). Following cold exposure with mild fasting, control mice showed modest reductions in adipose-related measures, whereas these changes tended to be attenuated after *Nts*+ SPN ablation, although the effects were small and did not consistently reach statistical significance. Notably, when food was freely available before and during cold exposure, females in the *Nts*+ SPN ablation group maintained both Tc and plasma NEFA levels (Figure S7). This nutritional rescue indicates that *Nts*+ SPNs become particularly important when cold defense must rely on endogenous energy reserves. Together, these loss-of-function results demonstrate that lower-thoracic *Nts*+ SPNs are required for efficient mobilization of stored lipids and maintenance of body challenge in the absence of food in female mice.

## Discussion

This study establishes a cell-type-resolved framework for understanding how the spinal sympathetic system converts physiological demands into selective peripheral outputs. By combining STx with activity mapping and circuit analysis, we identify the preganglionic layer as an organizing interface between central autonomic commands and organ-biased sympathetic control.

### Spatial and functional organization of SPNs

Our data suggest three principles of SPN organization. First, transcriptomic identity is coupled to spinal position. SPN subtypes showed characteristic rostrocaudal and mediolateral distributions, extending classical anatomical evidence for organotopic sympathetic organization^1,2^ to the molecular level. The male-biased enrichment of *Penk*+ and *Sst*+ SPNs in the upper lumbar cord further suggests their potential role in male-specific pelvic functions. More broadly, our atlas provides a roadmap for future studies linking SPN subtypes across the spinal cord to their upstream inputs, peripheral targets, and physiological functions.

Second, physiological challenges selectively recruit defined combinations of SPN subtypes rather than uniformly activating local sympathetic output. Cold and glucoprivation recruited partially overlapping but distinct populations (Figure 2), arguing against a purely global change in sympathetic tone^4^. In addition, spinal location within a transcriptomic subtype could bias recruitment, as illustrated by *Crh*+ SPNs. Thus, molecular identity and anatomical position together appear to shape functional recruitment.

Third, molecularly defined SPNs preferentially engage distinct peripheral pathways. Lower-thoracic *Cartpt*+, *Oxtr*+, and *Nts*+ SPNs show strongly biased projections toward the CG/SMG, adrenal medulla, and sympathetic trunk/ARG, respectively^12^ (Figure 3). Together with recent studies demonstrating molecular specialization among postganglionic sympathetic neurons^5–9^, these findings support a hierarchical modular organization of organ-selective sympathetic control across both pre- and postganglionic layers.

Several limitations should be emphasized. We examined only two physiological challenges, and it remains unknown how broadly the observed modular organization applies across sympathetic functions. The function of the cold- and glucoprivation-responsive *Crh*+ population remains unknown. IEG expression provides only an indirect and temporally limited measure of neuronal recruitment. In addition, current single-marker genetic approaches do not allow selective access to atlas-defined SPN subtypes. Our *Nts*-*Cre*-based manipulation includes multiple *Nts*-expressing populations and showed substantial ectopic viral labeling outside the IML (Figure S4 and Supplementary Note 1). Although anatomical, denervation, and ablation controls support a sympathetic efferent contribution to the observed phenotypes, more precise intersectional tools will be required to test the functions of individual transcriptomic SPN subtypes directly.

### Female-biased influence of *Nts*+ SPNs on WAT metabolism and cold defense

A notable feature of the *Nts*+ pathway is that its metabolic effects were strongly female-biased despite little detectable sex difference in the organization of the SPNs themselves. *Nts*+ SPN abundance, cold recruitment, axonal projections, and postganglionic activation were broadly similar between sexes (Figures 1, 3, and S3), whereas activation increased lipid mobilization and depleted WAT much more strongly in females, and ablation impaired cold-induced NEFA mobilization and cold tolerance selectively in females (Figures 5–7). This dissociation suggests that sex differences in sympathetic physiology can arise downstream of a largely shared preganglionic command.

Potential sites of this divergence include postganglionic neurons and adipose tissue. Estrogen signaling can enhance adipocyte lipolytic capacity and influence sympathetic regulation of WAT^43–46^, while sensory feedback from adipose tissue can restrain sympathetic and metabolic output^47,48^, providing additional potential sites at which sex-dependent modulation could arise. The relative contributions of these mechanisms and their underlying sexually dimorphic molecular and cellular processes remain elusive and warrant future investigation.

The fasting dependence of the loss-of-function phenotype suggests that *Nts*+ SPNs become important when cold defense relies on endogenous energy stores. Ablation produced no overt chronic adipose phenotype in the absence of cold exposure (Figure S6), and the thermal deficit was rescued when food remained available during cold exposure (Figure S7). This is consistent with previous evidence that WAT lipolysis becomes particularly important for cold tolerance when exogenous nutrients are limited^25^. One possibility is that females rely more strongly on lipid-fueled non-shivering thermogenesis^49–51^, whereas males may have greater capacity for muscle-based thermogenesis^52,53^. Future molecular and cellular studies are required to test these hypotheses and delineate the circuit-level and endocrine mechanisms underlying sex-specific thermoregulatory strategies.

In sum, our study connects SPN subtype organization, stressor-selective recruitment, peripheral pathway bias, organ regulation, and whole-body adaptation into a unified logic of sympathetic control. Extending this logic across organs, physiological states, and autonomic divisions may provide a general framework for deciphering how the nervous system orchestrates body-wide homeostasis.

## Supporting information

Soure Data

## Resource Availability

### Lead contact

Further information and requests for resources and reagents should be directed to and will be fulfilled by the Lead Contact, Kazunari Miyamichi.

### Materials availability

No new materials were generated in this study.

### Data and code availability

- STx data are available from the Gene Expression Omnibus (GEO) at GSE336196.
- All the other data generated in this study are presented in the main text or the supplemental materials.
- The custom scripts used to analyze the snRNA-seq and STx data in this study are available at https://github.com/gennkenn/Spatial-Transcriptomic-Activity-Atlas-of-Spinal-Sympathetic-Neurons.

## Acknowledgments

We thank Takeshi Sakurai for sharing *Nts-Cre* mice, RIKEN BDR laboratory for developmental genome system support with the Xenium and snRNA-seq analyses, RIKEN BDR laboratory for animal resources and genetic engineering for animal care and in vitro fertilization, Hinako Takase for technical assistance with FFPE preparation and HE staining, Chen Ran, Yuko Okamatsu, and members of the Miyamichi Laboratory for critical reading of the manuscript, and Addgene and the University of North Carolina Vector Core for AAV production. S.Y. is supported by a JSPS research fellowship for young scientists (DC), and H.S. by a JSPS postdoctoral research fellowship (PD). This work was supported by Japan Science and Technology Agency, CREST grant JPMJCR2021, Japan Society for the Promotion of Science, KAKENHI grant 23H04939, 23H04945, 25K21758, and 25K02368, the UEHARA Memorial Foundation, and RIKEN BDR Center Projects to KM.

## Author contributions

S.Y., H.S., and K.M. conceived the experiments. H.S. performed the STx experiments and analyzed the data with technical support by M.T., D.C., M.K., and T.K. S.Y. performed anatomical and functional characterization of SPNs with support by S.U. S.Y., H.S., and K.M. wrote the paper with contributions from all co-authors.

## STAR★Methods

### Key resources table

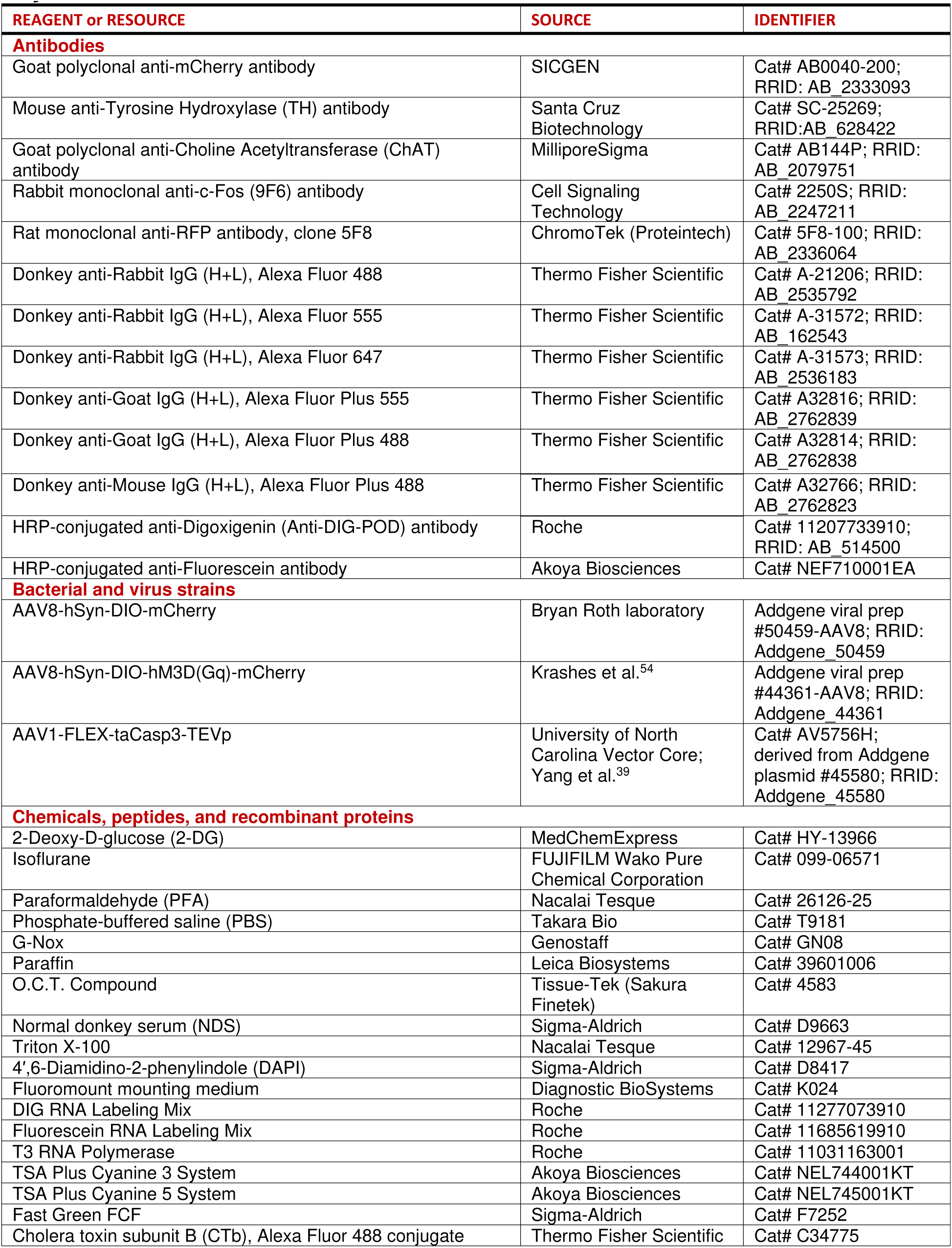
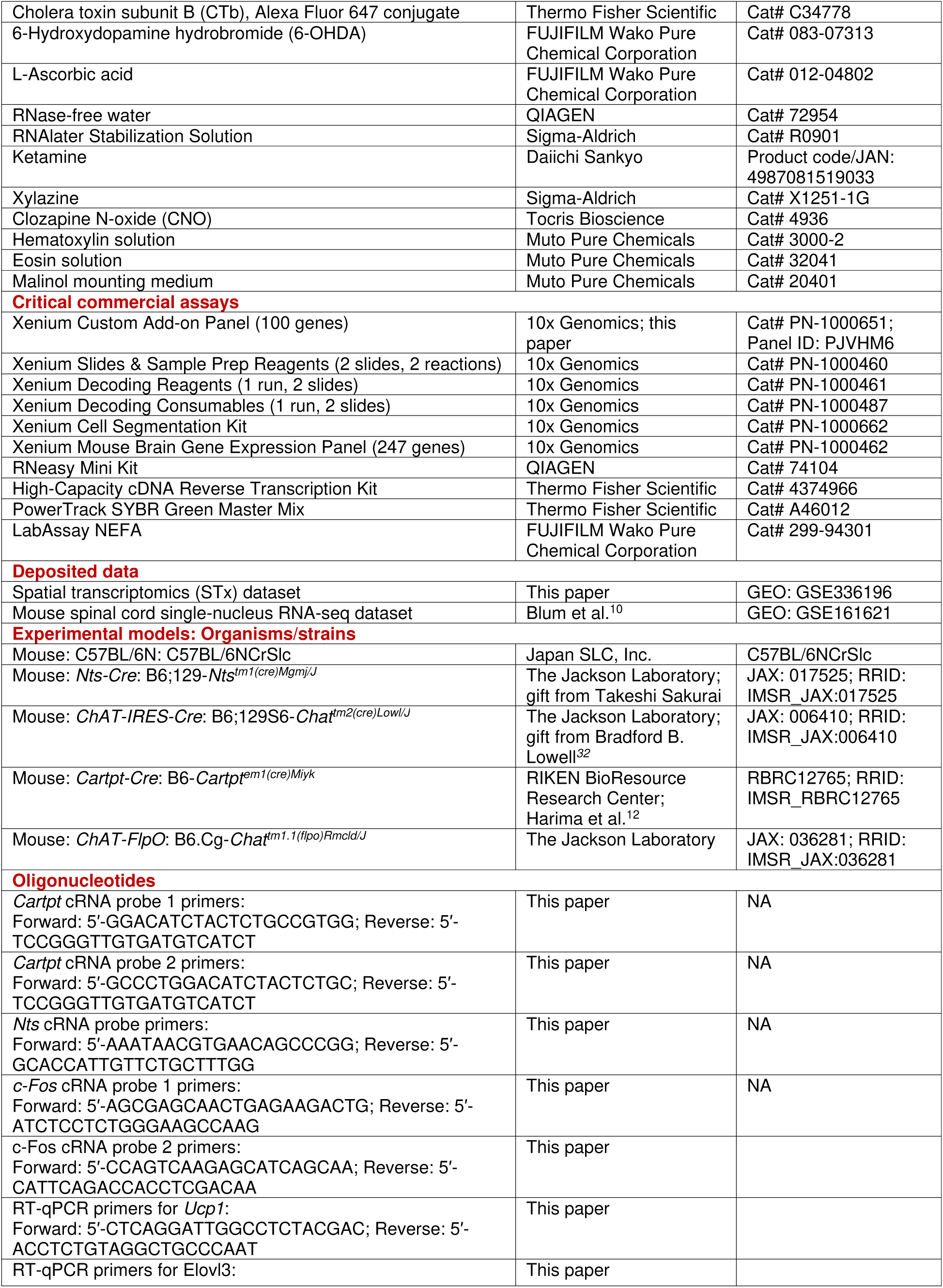
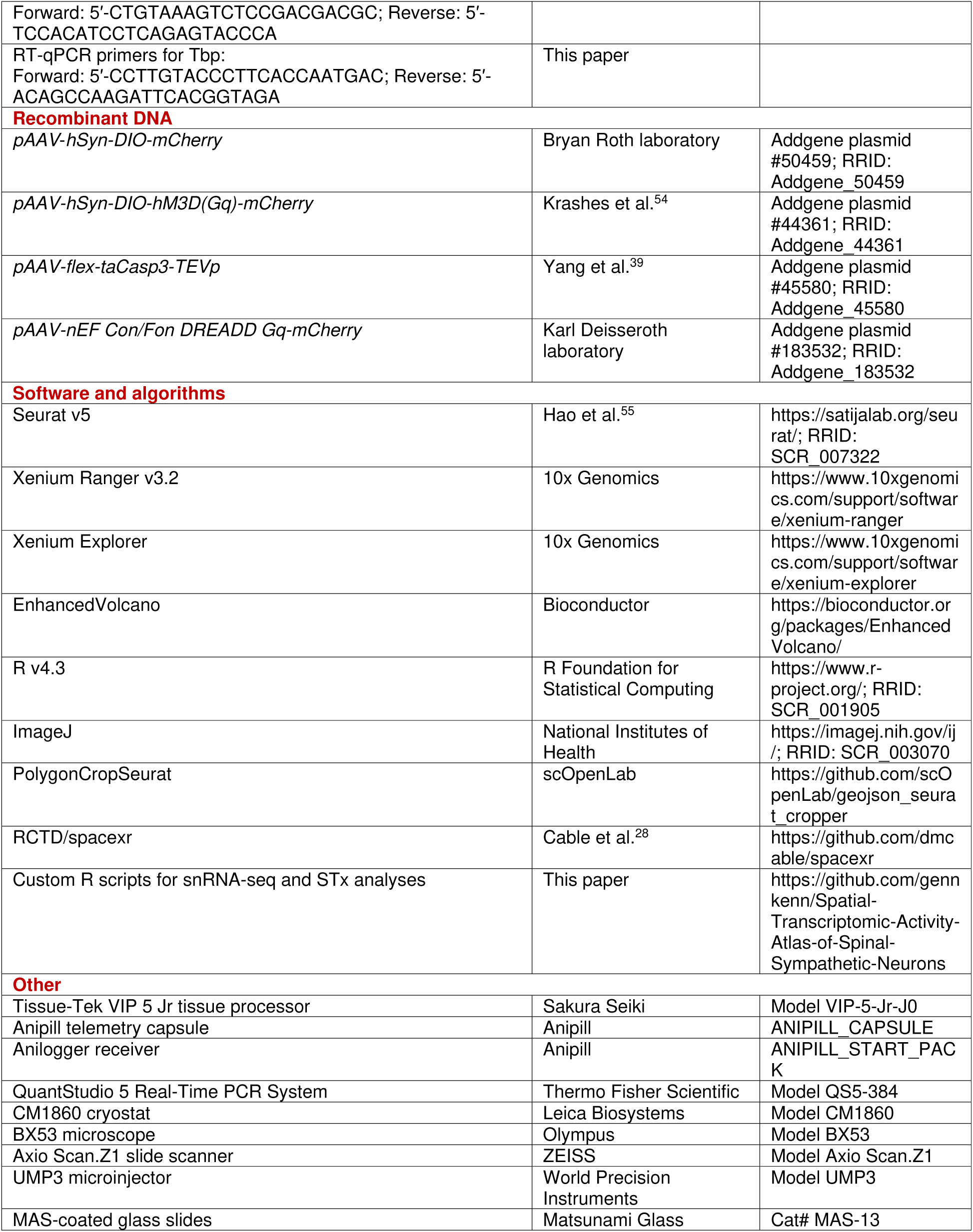

### EXPERIMENTAL MODEL AND SUBJECT DETAILS

All animal experiments were approved by the Institutional Animal Care and Use Committee of the RIKEN Kobe Campus. C57BL/6N mice were purchased from Japan SLC, Inc. (Shizuoka, Japan). *Nts-Cre*^31^ (Jax #017525) was a gift from Dr. Sakurai. *Chat-Cre*^32^ (Jax #006410) mice were a kind gift from B. Lowell. *Cartpt-Cre* mice have been described previously^12^. Animals were maintained at the animal facility of the RIKEN Center for Biosystems Dynamics Research under an ambient temperature of 21–24°C and a 12-hour light/12-hour dark cycle schedule. The mice were allowed ad libitum access to a laboratory diet (MFG; Oriental Yeast, Shiga, Japan; 3.57 kcal/g) and water, unless otherwise stated.

### METHOD DETAILS

#### Selection of Xenium custom probes

For the selection of Xenium custom probe sets, publicly available snRNA-seq data from Blum *et al.* (GSE161621)^10^ were analyzed. Six datasets were downloaded and processed in R v4.3 using the Seurat v5 package^55^. As an initial quality control step, nuclei with fewer than 400 or more than 8,000 detected genes, or with >10% of reads derived from mitochondrial or ribosomal genes, were excluded. The remaining nuclei were normalized using the *SCTransform* function^56^, regressing out both the mitochondrial and ribosomal gene ratios. Dimensionality reduction and integration were performed using *RunPCA*, *IntegratedLayers* (CCA-based integration), *RunUMAP*, *FindNeighbors*, and *FindClusters*, using the first 50 principal components (PCs) and a clustering resolution of 0.5. Clusters were annotated based on established marker genes for major cell types, including neurons (*Camk2a* and *Rbfox3*), mature oligodendrocytes (*Mog*), immature oligodendrocytes (*Pdgfra*), newly formed oligodendrocytes (*Bcas1*), endothelial cells (*Pecam1*), and astrocytes (*Aqp4*). Clusters expressing markers from multiple cell types or containing a low number of detected genes were excluded from further analyses.

For sub-clustering, neurons were independently reanalyzed. Clustering was performed using 50 PCs with a resolution of 0.8. Doublets and low-quality nuclei were removed based on the aforementioned criteria. Neuronal clusters were further classified into excitatory, inhibitory, and cholinergic neurons based on the expression of *Slc17a6*, *Gad1*, and *Slc5a7*.

Cholinergic neurons were subsequently sub-clustered using 50 PCs at a resolution of 0.5. Based on *Nos1* expression, sympathetic preganglionic neurons (SPNs) were identified and reanalyzed using 50 PCs at a resolution of 0.9. Clustering resolution was increased incrementally until no additional marker genes capable of distinguishing newly separated clusters could be identified.

To define marker genes for SPN clusters, skeletal motor neurons, and cholinergic interneurons, the *FindMarkers* function was applied using a significance threshold of *p* < 0.01 and an absolute log_2_ fold change > 0.5. Non-protein-coding genes were excluded from analysis. The final set of genes selected for Xenium custom probe design is listed in Table S1 and detailed data from this reanalysis are presented in Table S2.

#### Preparation of mouse spinal cord samples for the Xenium assay following stress conditions

Mice were individually housed for at least 7 days before exposure to the stress conditions. For each condition, N = 3 male and 3 female mice (10 weeks old) were used.

For the room temperature (RT) and saline control, mice received an intraperitoneal injection of saline (150 μL) and were maintained at room temperature for 1.5 h prior to tissue collection. For cold exposure, mice were placed in their home cages within a 4°C incubator for 1.5 h, followed immediately by perfusion. For the 2-DG condition, 2-DG (400 mg/kg; MCE, HY-13966) was intraperitoneally administered, and the mice were sacrificed 1.5 h later.

For tissue collection, mice were deeply anesthetized with an overdose of isoflurane and transcardially perfused with PBS, followed by 4% paraformaldehyde (PFA) in PBS. Tissues were post-fixed overnight in 4% PFA at 4°C and subsequently dehydrated in 70% ethanol at 4°C for at least one week. The thoracic, lumbar, and sacral spinal cord segments were dissected and subdivided into eight regions (T1–4, T5–7, T8– 10, T11–13, L1–3, L4–5, L6–S1, and S2–4) before dehydration. Dehydration and paraffin infiltration were conducted by Tissue-Tek VIP 5 Jr. (Sakura Seiki) using the settings as follows: 90 min in 80% ethanol, 90 min in 90% ethanol, 90 min in 100% ethanol three times, 90 min in 50% G-Nox/ethanol (Genostaff, GN08) three times, 90 min in G-Nox three times, and 90 min in paraffin (Leica, 39601006) four times. After paraffin infiltration, formalin-fixed, paraffin-embedded (FFPE) blocks were prepared.

FFPE blocks were sectioned at a 5 μm thickness and mounted onto slides according to the manufacturer’s protocol (10x Genomics, CG000578). Sections from all experimental conditions, including all three animals per condition, were placed on slides. The numbers of sections per condition, animal, and spinal segment are shown in Table S3.

Xenium analysis was performed using the pre-designed Xenium Mouse Brain Gene Expression Panel (247 genes; 10x Genomics, PN-1000462) in combination with a custom add-on panel (100 genes; 10x Genomics, PN-1000651) and reagents (10x Genomics, PN-1000460, 1000461, 1000487, and 1000662) according to the manufacturer’s protocols (CG000580, CG000749, and CG000584).

#### Data analysis of the Xenium assay

The output data from Xenium Explorer were processed using Xenium Ranger v3.2. The resulting datasets were individually imported into R v4.3, and the following analyses were performed separately for each sex and condition using the Seurat v5 package. The data were normalized using the *SCTransform* function. Dimensionality reduction and clustering were performed using *RunPCA*, *RunUMAP*, *FindNeighbors*, and *FindClusters*, with 50 PCs and a resolution of 1.0. Clusters were annotated based on canonical marker genes for neurons (*Camk2a* and *Rbfox3*), oligodendrocytes (*Opalin*), oligodendrocyte progenitor cells (OPCs) (*Pdgfra*), microglia (*Cd68*), meninges (*Dcn*), endothelial cells (*Pecam1*), ependymal cells (*Spag16*), and astrocytes (*Aqp4*). Clusters expressing markers for more than one major cell type were excluded from downstream analyses.

For neuronal subclustering, all neuronal clusters were independently reanalyzed using 50 PCs with a resolution of 0.5. Putative doublets were removed using the aforementioned criteria. Neurons were classified as excitatory, inhibitory, or cholinergic neurons based on the expression of *Slc17a6, Gad1,* and *Chat,* respectively. After cholinergic neurons from all datasets were merged, cholinergic neurons were further subclustered using 50 PCs at a resolution of 1.0 to dissect cholinergic interneurons identified by *Pax2*, motor neurons identified by *Bcl6*, and SPNs identified by *Nos1*. The SPNs were reanalyzed using 50 PCs. The clustering resolution was incrementally increased to 1.2 until no additional marker genes capable of distinguishing newly separated clusters were detected. Clusters with marker genes for non-SPN cell types were excluded from further analyses. The remaining cells were re-normalized and re-clustered. This procedure was repeated until no cluster exhibited strong expression of markers characteristic of non-target cell types. The correspondence between cell IDs and annotated cholinergic subtypes is summarized in Table S4. The marker genes for each cluster were identified using the *FindMarkers* function with the following thresholds: adjusted p-value < 0.05, minimum pct > 0.1, and log_2_ fold change > 0.25.

The coordinates of the central canal were manually annotated for each section to quantify the spatial relationship between SPNs and the central canal (Figure 1G, H). SPNs within the same section were identified using the *PolygonCropSeurat* function (https://github.com/scOpenLab/geojson_seurat_cropper), based on the corresponding manual annotation. The Euclidean distance between the centroid of each SPN and the center of the corresponding central canal was then calculated. To integrate the STx data with snRNA-seq datasets (Figure S1E, F), RCTD was applied following the Seurat v5 spatial integration workflow: https://satijalab.org/seurat/articles/seurat5_spatial_vignette_2. To assess *c-Fos* expression, SPNs with ≥5 *c-Fos* transcripts per cell were defined as *c-Fos*+. The IEG score was calculated using the *AddModuleScore* function in Seurat as a background-corrected expression score for *Fos*, *Fosb*, *Npas4*, *Nr4a1*, and *Egr1*. These IEGs were selected because they showed low basal expression and clear induction following stimulation (Figure S2A). SPNs with an IEG score greater than two standard deviations above the mean IEG score of all SPNs were classified as IEG+.

To create volcano plots, the EnhancedVolcano package in R (https://bioconductor.org/packages/release/bioc/html/EnhancedVolcano.html) was used.

#### Viral preparations

The following AAV vectors were purchased from Addgene. The titer is expressed as genomic particles (gp) per mL.

AAV serotype 8 *hSyn-DIO-mCherry* (2.0 × 10^13^ gp/mL, 50459-AAV8)

AAV serotype 8 *hSyn-DIO-Gq-mCherry* (2.1 × 10^13^ gp/mL, 44361-AAV8)

The following AAV vector was purchased from the University of North Carolina Viral Core. AAV serotype 1 *FLEX-taCasp3-TEVP* (1.2 × 10^12^ gp/mL)

#### Stereotaxic injection

For viral injections into the lower thoracic spinal cord, *Chat-Cre*, *Cartpt-Cre*, or *Nts-Cre* mice (4–5 weeks old) were anesthetized by intraperitoneal injection of ketamine (65 mg/kg; Daiichi Sankyo, 4987081519033) and xylazine (13 mg/kg; Sigma, X1251-1G). Under anesthesia, an incision was made over the thoracic region. The paraspinal muscles and dorsal vertebral elements were carefully removed by using fine forceps to expose the spinal cord. A pulled glass pipette connected to a microinjector (World Precision Instruments, UMP3) was used to inject 100 nL of AAV vector (AAV8 *hSyn-DIO-mCherry* or AAV8 *hSyn-DIO-hM3D(Gq)-mCherry*) or 150 nL of AAV1 *FLEX-taCasp3-TEVp*, each containing 0.1% Fast Green (Sigma, F7252), at 10 injection sites distributed bilaterally across the T8–T12 spinal segments (infusion rate: 80 nL/min).

For CTb injection into iWAT, gWAT, and kidneys (Figure 4), the mice were anesthetized as described above. A small incision was made in the abdominal region to expose the organs. Using a glass pipette connected to a microinjector, 500 nL of 0.1% CTb–Alexa Fluor 647 or CTb–Alexa Fluor 488 (Thermo Fisher Scientific, C34778 and C34775) was injected into six distinct sites per organ (infusion rate: 150 nL/min).

For chemical denervation of iWAT (Figure 6M–Q), 6-OHDA (Wako, 083-07313) was freshly prepared on ice by dissolving it in saline containing 0.1% ascorbic acid to a final concentration of 9 mg/mL. *Nts-Cre* mice injected with AAV8 *hSyn-DIO-hM3D(Gq)-mCherry* were anesthetized as described above. Bilateral incisions were made to expose iWAT, and 500 nL of the 6-OHDA solution was injected into each side at eight distinct sites. CNO administration was initiated one week after 6-OHDA treatment.

#### Histology and histochemistry

Mice were transcardially perfused with PBS, followed by 4% paraformaldehyde (PFA) in PBS. Tissues were post-fixed overnight in 4% PFA in PBS at 4°C, washed three times with PBS for 10–30 min each, and cryoprotected in 30% sucrose in PBS at 4°C for 24 h. Tissues were then trimmed, embedded in O.C.T. compound (Tissue-Tek, 4583), sectioned coronally at 30 μm using a cryostat (Leica, CM1860), and placed on MAS-coated glass slides (Matsunami, MAS-13). For antigen retrieval, sections were treated with Tris-Cl EDTA (pH 9.0) containing 1% normal donkey serum (NDS; Sigma, D9663) in PBST (PBS with 0.1% Triton X-100). After retrieval, sections were washed three times with PBS and blocked with 5% NDS in PBST for 30 min at 25 °C. The following primary antibodies were used in this study: goat anti-mCherry (1:350; SICGEN, AB0040-200), mouse anti-TH (1:500; Santa Cruz Biotechnology, sc-25269), goat anti-ChAT (1:150; Sigma, AB144P), and rabbit anti-c-Fos (1:500; Cell Signaling Technology, 2250S). These sections were then washed three times with PBS and treated with the following secondary antibodies, diluted in 1% NDS in PBST and containing 4′,6-diamidino-2-phenylindole dihydrochloride (DAPI), for 2 h at room temperature or overnight at 4°C: donkey anti-rabbit Alexa Fluor 555/488/647 (1:300; Thermo Fisher Scientific, A31572, A21206, A31573), donkey anti-goat Alexa Fluor 555 (1:300; Thermo Fisher Scientific, A32816), and donkey anti-mouse Alexa Fluor 488 (1:500; Thermo Fisher Scientific, A32766). The sections were washed once with PBS and mounted under coverslips using Fluoromount (Diagnostic BioSystems, K024).

In situ hybridization (ISH) was performed as described previously^57^. To generate cRNA probes, DNA templates were amplified from mouse spinal cord cDNA by PCR (Genostaff, MD-23). A T3 RNA polymerase recognition site (5′-AATTAACCCTCACTAAAGGG) was added to the 3′ end of the reverse primers. The primer sets and sequences of the probe targets were as follows.

- *Cartpt*−1: 5′-ggacatctactctgccgtgg; 5′-tccgggttgtgatgtcatct
- *Cartpt*−2: 5′-gccctggacatctactctgc; 5′-tccgggttgtgatgtcatct
- *Nts*: 5′-aaataacgtgaacagcccgg; 5′-gcaccattgttctgctttgg
- *c-Fos*−*1*: 5′-agcgagcaactgagaagactg; 5′-atctcctctgggaagccaag
- *c-Fos*−*2*: 5′-ccagtcaagagcatcagcaa; 5′-cattcagaccacctcgacaa

PCR-amplified DNA templates (800 ng) were used for in vitro transcription with either DIG (Roche, 11277073910) or Flu (Roche, 11685619910) RNA-labeling mix and T3 RNA polymerase (Roche, 11031163001), following the manufacturer’s instructions (Roche). For *Cartpt* and *c-Fos* detection, the two probes were combined to improve the signal-to-noise ratio.

For single-color ISH combined with immunostaining, following hybridization and post-hybridization washes, tissue sections were incubated overnight with HRP-conjugated anti-DIG antibody (1:500; Roche, 11207733910), together with either goat anti-mCherry antibody (1:350; SICGEN, AB0040-200) or rat anti-RFP antibody (1:250; Proteintech Group, 5F8-100). Signal amplification was performed using TSA-plus Cyanine 3 (1:70; AKOYA Biosciences, NEL744001KT) for 25 min, followed by washing. mCherry-positive cells were visualized using a donkey anti-goat Alexa Fluor 488 antibody (1:200; Thermo Fisher Scientific, A32814). Nuclear counterstaining was carried out with PBS containing 50 ng/ml DAPI (Sigma-Aldrich, D8417).

For dual-color ISH combined with goat anti-ChAT immunostaining, Flu-labeled RNA probes were first detected using an HRP-conjugated anti-Flu antibody (1:250; AKOYA Biosciences, NEF710001EA), followed by signal amplification with TSA plus cyanine 3 (1:70 in 1× Plus Amplification Diluent). HRP activity was then quenched by incubation in 2% sodium azide in PBS for 15 min at 25 °C. Subsequently, DIG-labeled cRNA probes were detected using an HRP-conjugated anti-DIG antibody (1:500; Roche, 11207733910) and TSA-plus Cyanine 5 (1:70; AKOYA Biosciences, NEL745001KT). Finally, the sections were mounted using Fluoromount and covered with glass coverslips.

For hematoxylin and eosin (H&E) staining, adipose tissues (iWAT and gWAT) were fixed in 4% PFA in PBS for two nights at 4°C, followed by cryoprotection in 30% sucrose in PBS for two additional nights at 4°C. Tissues were then embedded in O.C.T. compound (Tissue-Tek, 4583) and sectioned at 15 μm thickness using a cryostat (Leica, CM1860). Frozen sections mounted on glass slides were air-dried for approximately 4 h, rinsed with distilled water, stained with hematoxylin (Muto Pure Chemicals, 3000-2) for 90 s, and washed under running tap water for 20 min. Sections were then counterstained with 1% eosin (Muto Pure Chemicals, 32041) diluted in 95% ethanol for 90 s. After staining, sections were dehydrated sequentially in 70% ethanol (1 min), 85% ethanol (1 min), 95% ethanol (1 min), and 100% ethanol (two changes, 1 min each), cleared in xylene (four changes, 1 min each), and mounted using malinol mounting medium (Muto Pure Chemicals, 20401).

#### Quantification of *in situ* hybridization and immunohistochemistry data

To quantify *c-Fos*+ SPNs within the IML and IC (Figures 3 and S3), T8–T12 coronal spinal cord sections were subjected to *in situ* hybridization using a DIG-labeled probe for *Nts* and fluorescein-labeled probes for *c-Fos*, followed by anti-ChAT immunostaining as described in the **Histology and histochemistry** section. Among ChAT+ neurons, cells positive for *Nts* and/or *c-Fos* were quantified using ImageJ software (NIH). Positive cells were identified and counted manually, and the fractions of *Nts*+ or *c-Fos*+ cells among ChAT+ neurons were calculated. For each animal, the values were averaged across at least ten sections, and the resulting mean was used for statistical analysis.

To quantify SPN-derived axons (Figures 3 and S3D–F), 30-μm-thick sections of the ST, ARG, CG/SMG, and adrenal medulla were immunostained with anti-TH and anti-mCherry antibodies as described in the **Histology and histochemistry** section. Images were acquired using an Olympus BX53 microscope equipped with a 10× objective (NA 0.4) for the CG/SMG and adrenal medulla and a 20× objective (NA 0.75) for the ST and ARG. For each image, the TH-positive area was manually delineated using ImageJ software. The mCherry-positive signal was binarized using the *Threshold* function in ImageJ, and the proportion of the mCherry-positive areas within the TH-positive region was calculated manually. For each animal, values were averaged from at least three images per section.

To quantify c-Fos+ and CTb+ cells (Figures 3, 4), 30-μm-thick sections of the CG/SMG, ST and ARG were immunostained with anti-c-Fos and anti-TH antibodies, as described above. The numbers of c-Fos+, TH+, CTb+, and DAPI+ cells were counted manually using ImageJ software. For each mouse, the ratio of c-Fos+ cells to TH+ neurons was calculated by averaging the values from at least three sections per ganglion. In Figure S5D, images were acquired using a Zeiss Axio Scan.Z1 with a 10× (NA 0.45) objective lens. The number of c-Fos+ cells was manually counted using ImageJ software. For each mouse, the cells were quantified from four sections per nucleus (two sections per hemisphere), with image brightness and contrast adjusted uniformly across all samples.

To quantify adipocyte size, images were acquired using an Olympus BX53 microscope equipped with a 10× objective lens (NA 0.4). Individual adipocytes were manually outlined in ImageJ, and cell size was calculated for each traced cell. For each animal, the measurements were averaged across at least three independent images, and the resulting mean value was used for statistical analysis.

#### Adipose RNA extraction and RT-qPCR analysis

To stabilize RNA, freshly collected adipose tissues were immersed in RNAlater (Sigma, R0901) and stored at −80 °C until processing. Total RNA was extracted from frozen tissues using the RNeasy Mini Kit (Qiagen, 74104) according to the manufacturer’s instructions. The RNA concentration was measured, and all samples were adjusted to 300 ng/μL with RNase-free water. Reverse transcription was performed using the High-Capacity cDNA Reverse Transcription Kit (Thermo Fisher Scientific, 4374966).

Quantitative real-time PCR (RT-qPCR) was performed by mixing cDNA templates with gene-specific primers and PowerTrack SYBR Green Master Mix (Thermo Fisher Scientific, A46012), followed by analysis using a QuantStudio 5 Real-Time PCR System (Thermo Fisher Scientific, QS5-384). Relative mRNA expression levels were calculated using the ΔΔCt method, with Tata-box binding protein (*Tbp*) used as the internal reference gene. Fold changes were calculated by determining the difference between each ΔCt and the mean ΔCt of the control group.

Primer sequences (5’-3’) were as follows:

*Ucp1*: ctcaggattggcctctacgac; acctctgtaggctgcccaat
*Elovl3*: ctgtaaagtctccgacgacgc; tccacatcctcagagtaccca
*Tbp*: ccttgtacccttcaccaatgac; acagccaagattcacggtaga

#### Measurement of plasma NEFA

Blood samples were collected from the facial vein and placed in ethylenediaminetetraacetic acid-coated tubes. Plasma was obtained by centrifugation, and NEFA concentrations were measured using a LabAssay NEFA kit (FUJIFILM Wako Pure Chemical Corporation, 299-94301), according to the manufacturer’s instructions.

#### Measurement of Tc

To continuously monitor Tc, Anipill telemetry capsules (Anipill) were surgically implanted into mice under anesthesia induced by intraperitoneal injection of a ketamine/xylazine mixture, as mentioned above. After surgery, the mice were allowed to recover for at least one week before the experiments. Tc data were recorded wirelessly using an Anilogger receiver (Anipill) placed near the home cage.

#### Chemogenetic manipulation and cell ablation

For chemogenetic activation of *Nts*+ or *Cartpt*+ SPNs, AAV8 *hSyn-DIO-Gq-mCherry* was injected into the spinal cord at the T8–T12 levels in *Nts-Cre*, *Cartpt*-*Cre*, or *Chat*-*Cre* mice, respectively. Experiments were initiated at least two weeks after surgery. For the RT-qPCR assay (Figures 5E–H), mice received intraperitoneal injections of CNO (Tocris Bioscience, 4936; 2 mg/kg) three times within a single day at 4-hour intervals (10:00, 14:00, and 18:00; Zeitgeber time 0 = 8:00). For the chronic chemogenetic activation experiments (Figure 6A–L), activation was initiated more than 2 weeks after viral injection, once the body weight of the mice reached 19–21 g. To activate *Nts*+ SPNs, mice were intraperitoneally administered CNO (2 mg/kg) once daily between 18:00 and 20:00 for seven consecutive days. For the cell ablation experiments (Figure 7), AAV1 *FLEX-taCasp3-TEVP* was injected into the T8–T12 segments of the spinal cord in *Nts-Cre* mice. Cold exposure experiments were conducted three weeks after surgery to allow sufficient time for cell ablation.

#### Quantification and statistics

Statistical analyses were performed using custom R scripts. All tests were two-tailed. The sample sizes and statistical tests are indicated in the figure and legends. Error bars are defined in figure legends.

## Supplementary Materials

Supplementary Text

Figures S1 to S7

Tables S1 to S4

References

## Supplementary Text

### Supplementary Note 1

Although the *Nts-Cre* mouse line enabled relatively restricted AAV targeting to *Nts*+ neurons in the IML (Figure S4A), where most *Nts*+ SPNs reside (Figure S3A), we observed substantial ectopic and nonspecific viral labeling outside the IML. In particular, putative sensory neurons in the dorsal horn along the injection tract were prominently labeled, regardless of *Nts* expression (Figure S4D). Because these non-ChAT+ spinal cells are unlikely to project to the sympathetic trunk, prevertebral ganglia, or adrenal medulla, the axonal mapping results shown in Figure 3 should reflect *Nts*+ SPNs. However, such nonspecific labeling can compromise the specificity of the manipulation experiments in Figures 5–7.

To reduce off-target labeling, we attempted an intersectional strategy using the *ChAT-FlpO* mouse line (JAX #036281), crossed this line with *Nts-Cre* to generate double-heterozygous animals, and injected custom AAV9 (*pAAV-nEF Con/Fon Gq-mCherry*; Addgene #183532). However, this approach yielded only a very small number of labeled cells. Given the low efficiency of the Con/Fon strategy, we proceeded with *Nts-Cre* alone and implemented the following controls to aid in interpretation.

For gain-of-function experiments (Figures 5 and 6), we chemically denervated iWAT using 6-OHDA (Figure 6M–Q), which reversed the lipolytic phenotype induced by activation of hM3Dq+ cells in *Nts-Cre* mice. This suggests that lipolysis is mediated by sympathetic activation of WAT-innervating postganglionic neurons rather than by systemic effects. For loss-of-function experiments (Figure 7), we confirmed a substantial reduction of *Nts*+ SPNs in the IML without a detectable reduction of *Nts*+ sensory neurons in the dorsal horn and that central cold-sensing and thermogenic neurons remained functionally intact following ablation in *Nts-Cre* mice (Figure S5). Accordingly, we interpreted the observed effects on thermoregulation and NEFA levels as arising from the loss of sympathetic efferent function rather than from sensory or central deficits.

## Supplementary Figure

**Figure S1:**
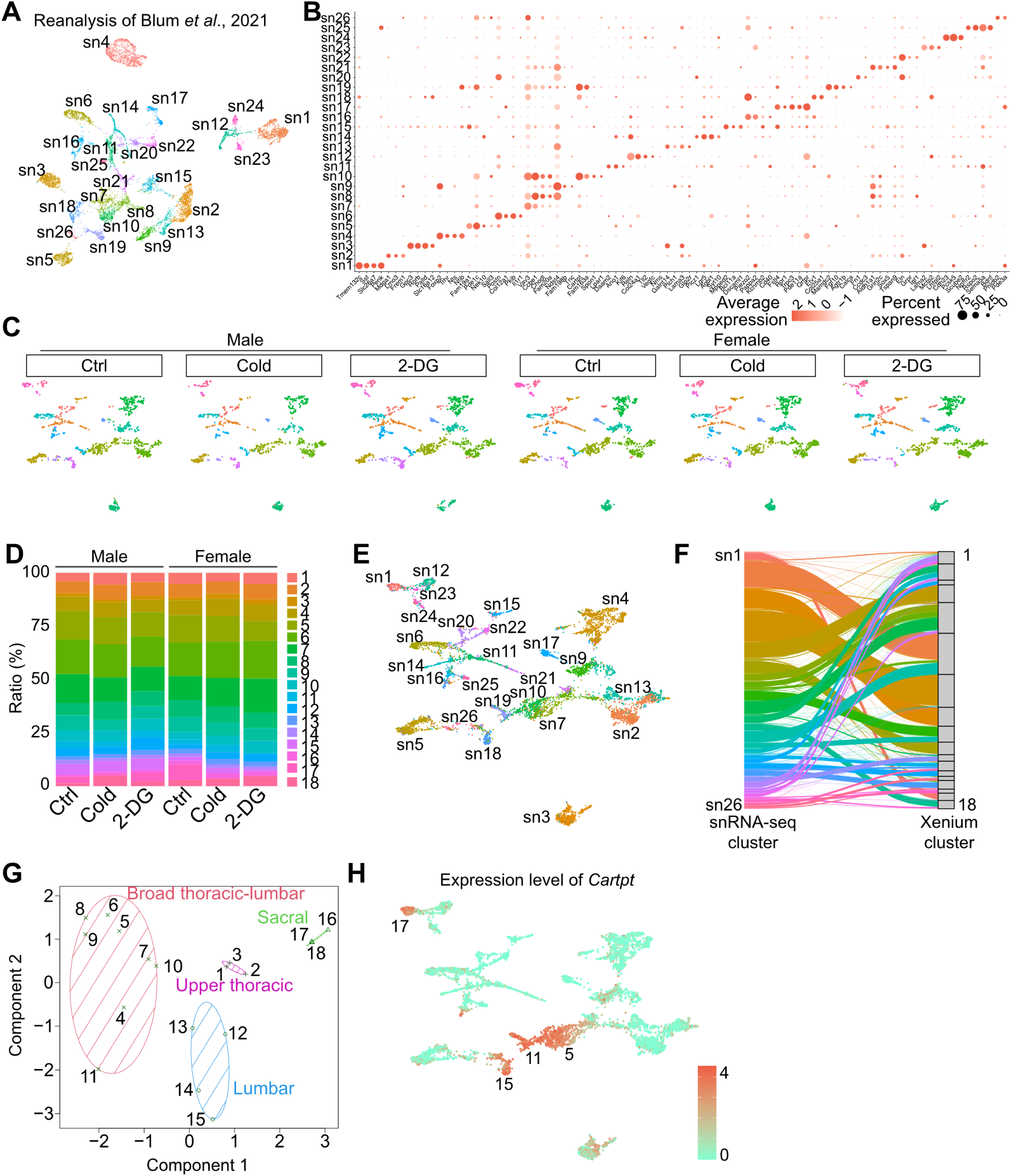
Reanalysis of published SPN snRNA-seq data for Xenium custom probe design and additional STx analyses, related to Figure 1. (A) UMAP representation of reanalyzed snRNA-seq data from Blum *et al*.^10^. After quality control, transcriptomic profiles were obtained from 8,553 SPNs and classified into 26 clusters. The prefix “sn” indicates cell types identified in the snRNA-seq dataset. (B) Dot plot showing marker genes for the clusters defined in (A). (C) UMAP representations of STx data for male (left) and female (right) SPNs, separated by experimental condition. Dot colors indicate cell types classified using canonical marker genes^10^. (D) Bar graphs showing the proportion of each SPN subtype, demonstrating comparable cluster distributions across conditions in both sexes. (E, F) UMAP (E), and Sankey plot (F) showing the correspondence between cell types in the snRNA-seq dataset, indicated by the prefix “sn,” and clusters in the STx dataset, as predicted by RCTD^28^. (G) *k*-means clustering of SPN subtypes based on their rostrocaudal distribution, identifying four spatial distribution patterns corresponding to those defined in Figure 1E, F. (H) UMAP representation showing *Cartpt* expression in SPNs. Darker red indicates higher expression. Prominent expression was found in clusters 5, 11, and 15. For more data, see Table S2 and Source Data file.

**Figure S2:**
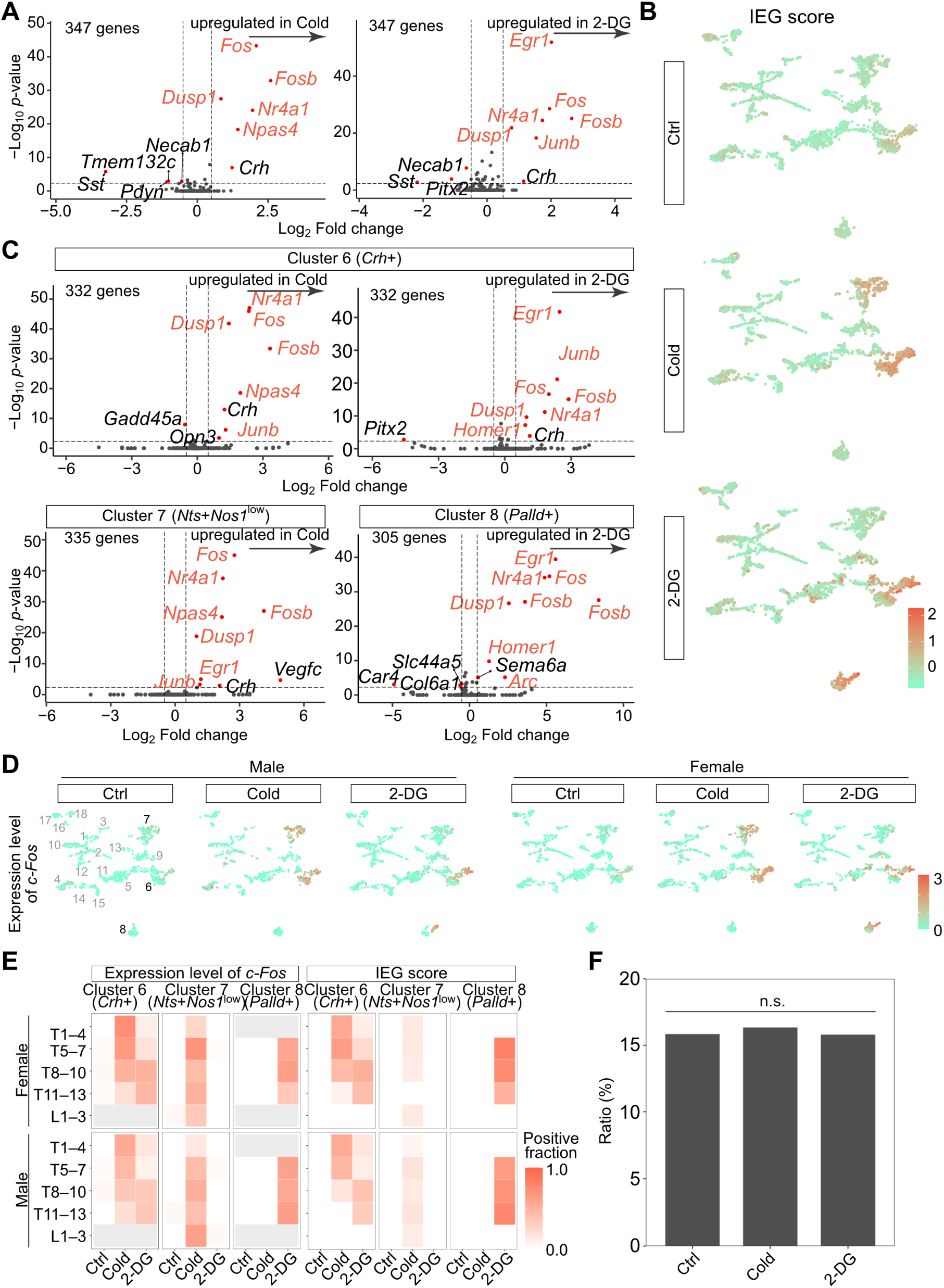
Additional analyses of IEG expression, related to Figure 2. (A, C) Volcano plots comparing gene expression in pan-SPNs (A) or in specific clusters (C) between control and cold exposure conditions (left) or between control and 2-DG conditions (right). The x-axis shows log_2_-transformed fold change, and the y-axis shows −log_10_-transformed *p*-values from the Wilcoxon rank-sum test. Genes with a fold change >2 or <0.5 and *p* < 0.01 were defined as differentially expressed genes (DEGs; red dots). IEGs among the DEGs are indicated in red text. (B) UMAP representations of IEG scores across the three conditions. A darker red color indicates higher expression. (D) UMAP representations of IEG scores across the three conditions for each sex. (E) Heatmaps showing the *c-Fos*+ (left) or IEG+ (right) fractions in each condition across the thoracolumbar segments for each sex. Segments accounting for less than 5% of all the SPNs in each subtype were excluded from the display. (F) Bar graph showing the fraction of cluster 6 (*Crh*+ SPNs) under each condition. Fisher’s exact test with Holm’s correction: n.s., not significant. These data indicate that although *Crh* expression is upregulated by cold exposure and 2-DG administration (A, C), it does not affect the overall classification of transcriptomic SPN subtypes. For more data, see Source Data file.

**Figure S3:**
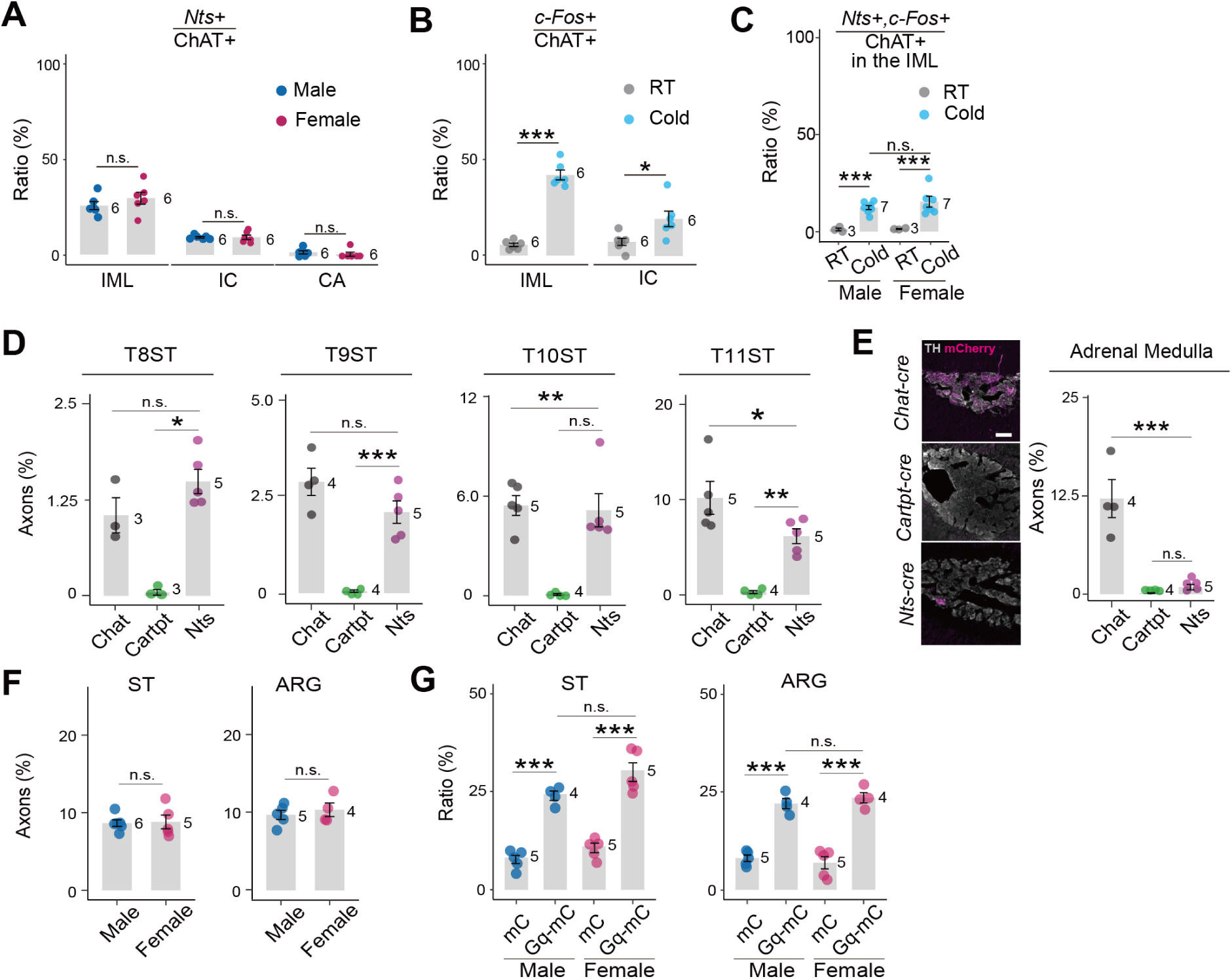
Additional characterization of *Nts*+ SPN identity and axonal projections, related to Figure 3. (A) Fraction of *Nts*+ SPNs among ChAT+ cells in the IML, IC, and CA regions of male and female mice. (B) Fraction of *c-Fos*+ cells among the ChAT+ cells in the IML (left) and IC (right) at RT and after cold exposure. In (A) and (B), n.s., not significant; \**p* < 0.05; \*\*\**p* < 0.001 by two-sided Welch’s t-test. (C) Fraction of *Nts*+, *c-Fos*+ cells among ChAT+ cells in the IML at RT and after cold exposure in male and female mice. One-way ANOVA followed by Tukey’s post hoc test: n.s., not significant; \*\*\**p* < 0.001. (D, E) Quantification of mCherry+ axons originating from *Chat+*, *Cartpt+*, and *Nts+* SPNs in the lower ST (T8–T11) (D) and adrenal medulla (E), as shown in Figure 3E. Representative images of the adrenal medulla are shown on the left in (E). One-way ANOVA followed by Tukey’s post hoc test: \**p* < 0.05, \*\**p* < 0.01, \*\*\**p* < 0.001. (F) Quantification of mCherry+ axonal areas within the TH+ regions originating from *Nts*+ SPNs in the ST (left) and ARG (right) of male and female mice. n.s., not significant using two-sided Welch’s t-test. (G) Fraction of c-Fos+ cells among TH+ cells in the ST (left) and ARG (right) following chemogenetic activation of *Nts*+ SPNs in male and female mice, corresponding to Figure 3G. One-way ANOVA followed by Tukey’s post hoc test: \*\*\**p* < 0.001. The number of animals in each group is shown. Error bars represent the standard deviation.

**Figure S4:**
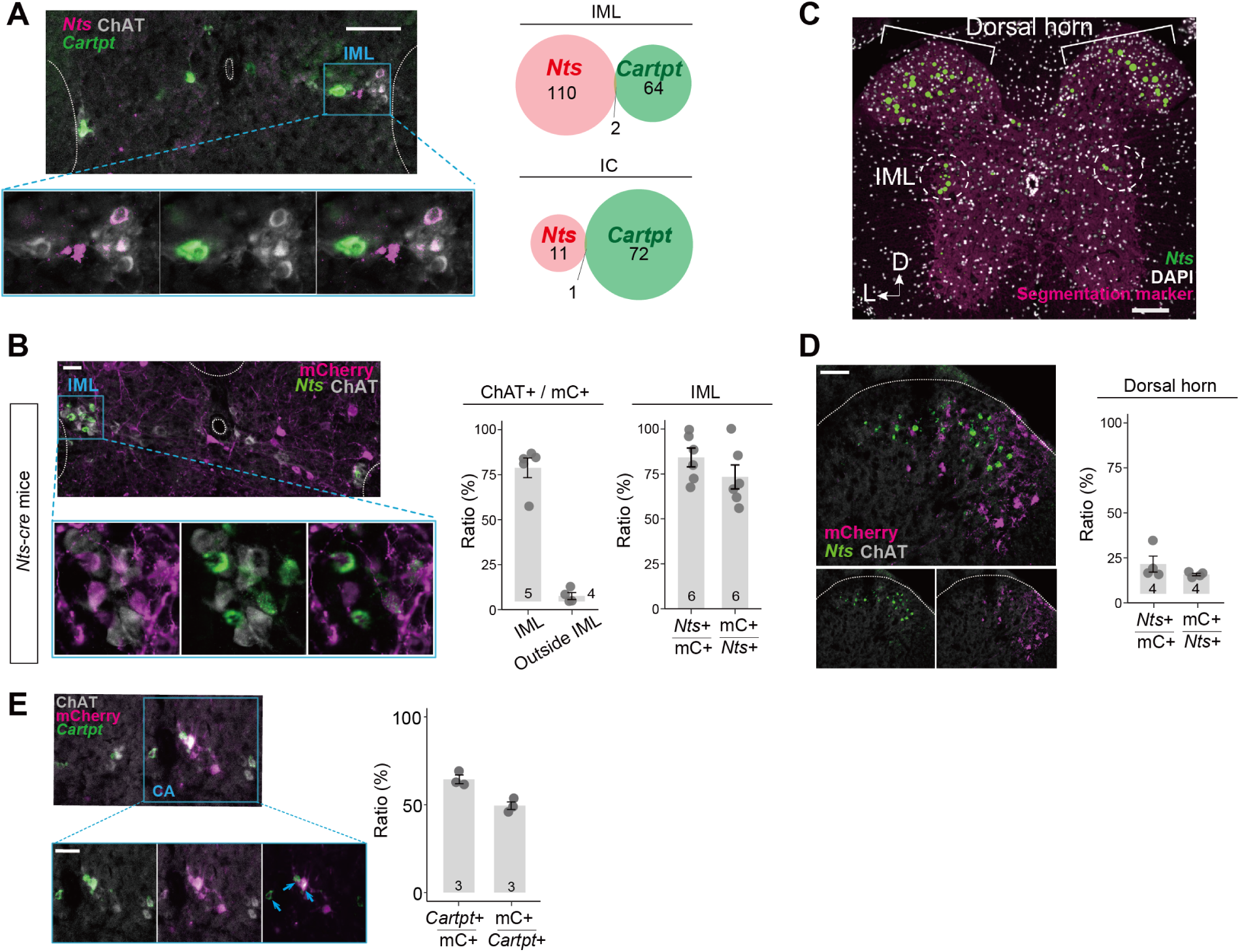
Characterization of *Nts*-Cre- and *Cartpt*-Cre-based viral targeting, related to Figures 3 and 5. (A) Left: Representative spinal cord section showing *Cartpt* (green), *Nts* (magenta), and ChAT (gray). Right: Quantification of *Cartpt*+, *Nts*+, and double-positive cells in the IML (top) and IC (bottom), showing that *Cartpt*+ and *Nts*+ SPNs were largely non-overlapping. Data were pooled from N = 4 wild type mice. (B) Left: Representative spinal cord section from *Nts-Cre* mice used in Figure 3D, E, showing *Nts* (green), mCherry (mC, magenta), and ChAT (gray). Middle: Fraction of ChAT+/mC+ cells within and outside the IML. Viral labeling within the IML predominantly occurred in ChAT+ neurons, whereas most mC+ cells outside the IML were ChAT−, indicating ectopic and non-specific AAV targeting. Right: Quantification of AAV injection efficiency (mC+/*Nts*+) and specificity (*Nts*+/mC+) within the IML. (C) Representative image of the T8–T10 spinal cord from the STx dataset showing *Nts* expression (green circles, with circle size indicating expression level), segmentation markers (ATP1A1, E-cadherin, and CD45; magenta), and DAPI staining (light gray). D, dorsal; L, lateral. (D) Left: Representative spinal cord section from *Nts*-Cre mice showing *Nts* (green), mCherry (magenta), and ChAT (gray) in the dorsal horn. Right: Quantification of the AAV targeting efficiency (mC+/*Nts*+) and specificity (*Nts*+/mC+) within the dorsal horn. These data show substantial ectopic viral labeling in this region. (E) Validation of hM3Dq-mCherry targeting in *Cartpt-Cre* mice used for the chemogenetic activation experiment in Figure 5D. Left: Representative spinal cord section showing *Cartpt* (green), mCherry (magenta), and ChAT (gray) in the central autonomic area. Right: Quantification of viral targeting specificity (*Cartpt*+/mC*+*) and efficiency (mC*+*/*Cartpt*+). Blue arrows indicate representative *Cartpt*+ mCherry+ cells. Scale bars, 100 μm in (A and C) and 50 μm in (B, D, and E). The number of animals in each group is shown. Error bars represent the standard deviation. See Supplementary Note 1 for further interpretation.

**Figure S5:**
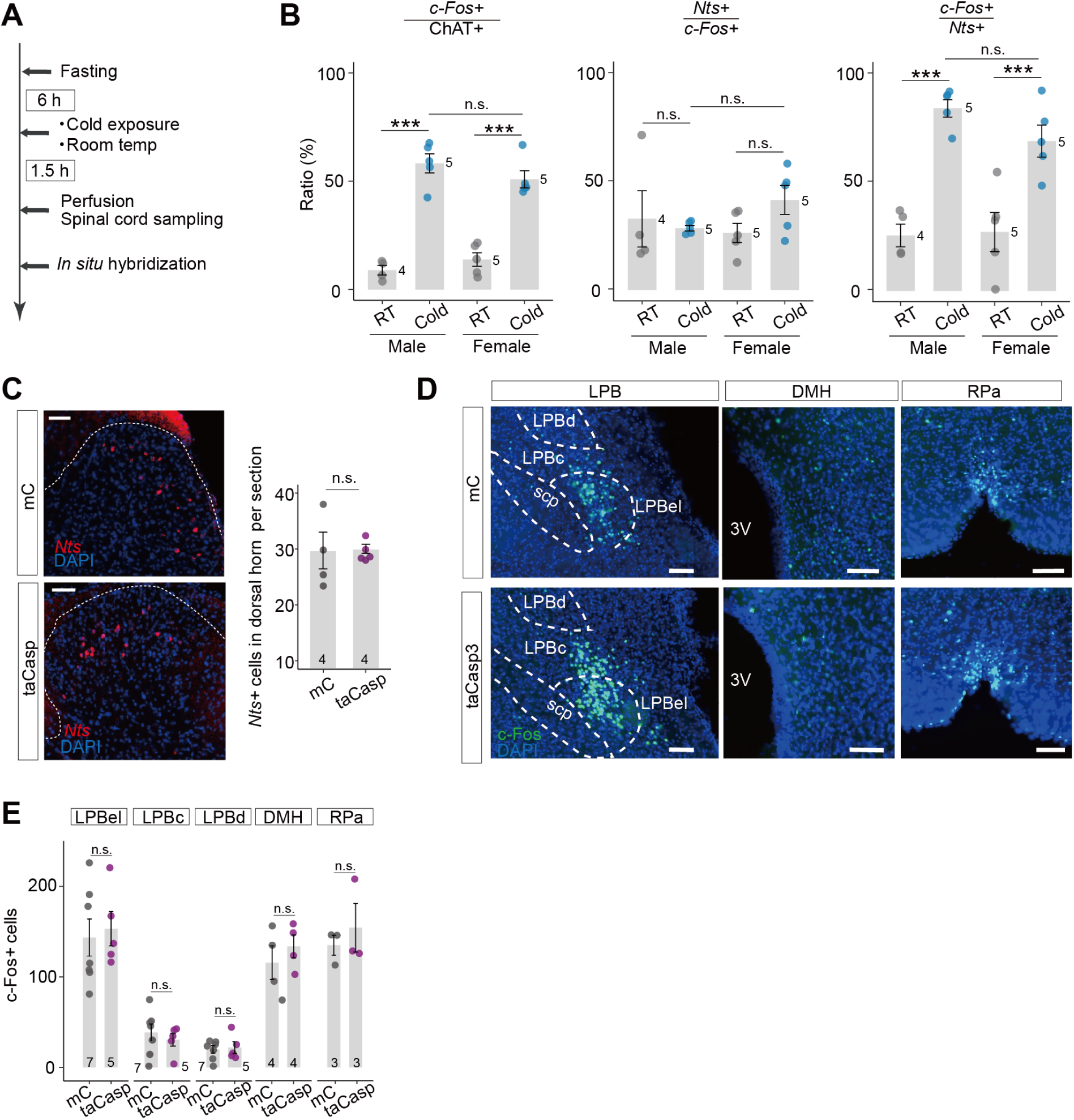
The *Nts*+ SPN ablation procedure does not affect the cold sensory system, related to Figure 7. (A) Experimental scheme and timeline for the *c-Fos* expression assay after cold exposure under conditions without food access, as shown in Figure 7A. (B) Quantification of the indicated parameters in both sexes. One-way ANOVA followed by Tukey’s post hoc test: \*\**p* < 0.01, \*\*\**p* < 0.001. Compared to Figure 3C, these data obtained under mild fasting conditions reveal: i) slightly higher basal *c-Fos* levels in RT controls; and ii) comparable activation of *Nts*+ SPNs in both sexes. (C–E) Sensory components were analyzed according to the scheme shown in Figure 7A. Some of the animals were also used in the experiments shown in Figure 7. (C) Left: Coronal spinal cord sections showing *Nts*+ sensory neurons (red) and DAPI (blue) in the dorsal horn of control (mC; left) and taCasp (right) mice. Right: Quantification of *Nts*+ sensory neurons per section in the dorsal horn of the control and taCasp groups. n.s., not significant according to a two-sided Welch’s t-test. (D) Coronal brain sections, including the lateral parabrachial nucleus (LPB), dorsomedial hypothalamus (DMH), and raphe pallidus nucleus of the medulla (RPa), showing c-Fos (green) and DAPI (blue) staining in mC and taCasp mice. LPB subdivisions: d, dorsal; c, central; el, external lateral. scp, superior cerebellar peduncle. (E) Quantification of *c-Fos*+ neurons in each brain area in the mC and taCasp groups. n.s., not significant according to two-sided Welch’s t-test. These data suggest that the cold sensory system remained intact in the *Nts*+ SPN ablation group. The number of animals in each group is shown. Error bars represent the standard deviation. Scale bar, 100 μm.

**Figure S6:**
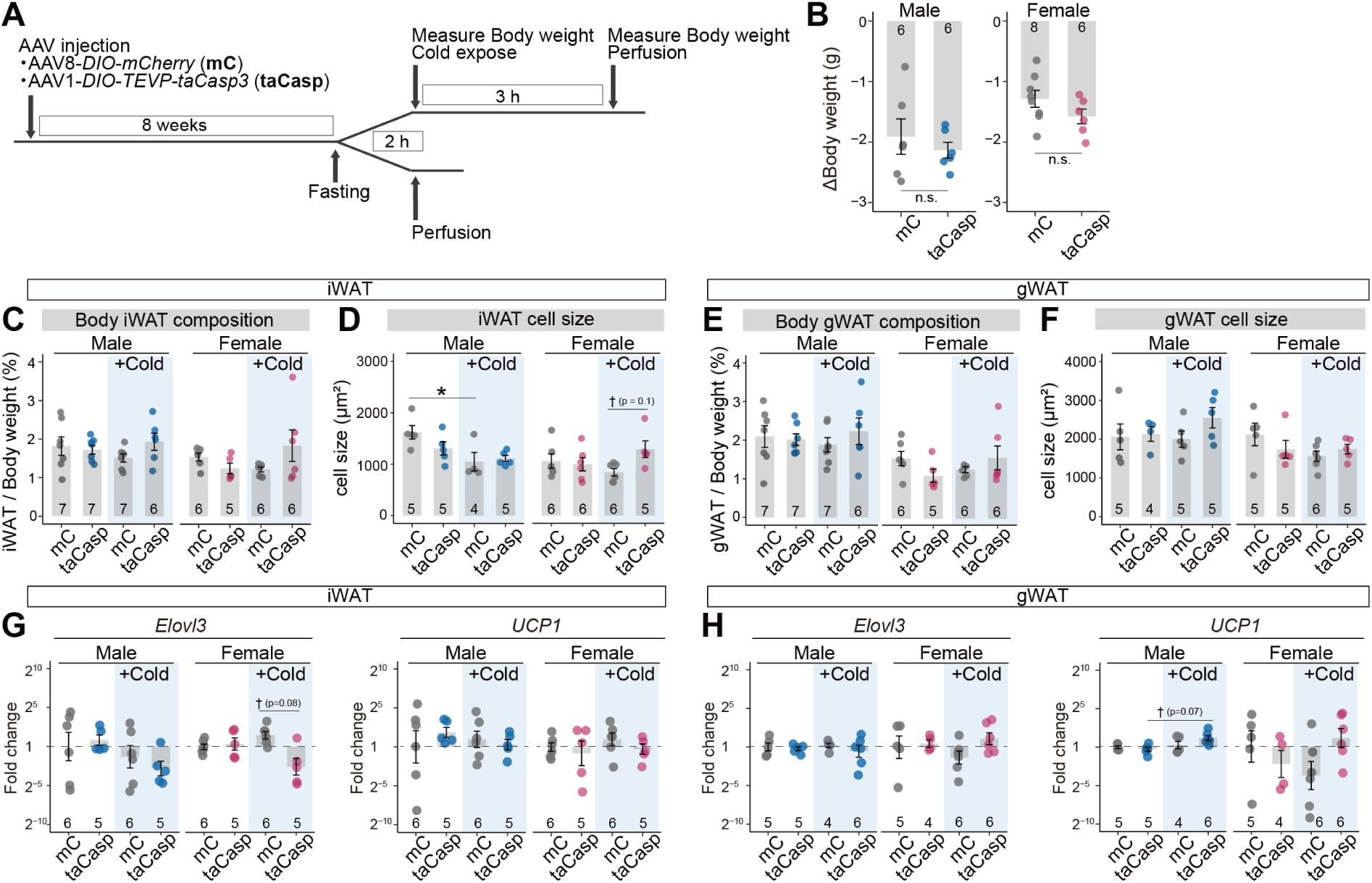
Effects of chronic *Nts*+ SPN ablation on adipose tissue under cold challenge when food is unavailable, related to Figure 7. (A) Experimental scheme for measuring the effects of chronic *Nts*+ SPN ablation on adipose tissue with or without cold exposure. (B) Change in body weight during the cold-challenge period in control (mC) and taCasp mice. (C, E) Relative mass of iWAT (C) and gWAT (E) in male and female control and taCasp mice with or without cold exposure. (D, F) Quantification of adipocyte size in iWAT (D) and gWAT (F) under the same conditions. (G, H) Fold changes in *Elovl3* and *Ucp1* expression in iWAT (G) and gWAT (H) from male and female control and taCasp mice with or without cold exposure. One-way ANOVA with post hoc Tukey’s test: n.s., not significant; \**p* < 0.05; Exact p values are indicated for comparisions marked with †. The number of animals in each group is shown. Error bars represent the standard deviation.

**Figure S7:**
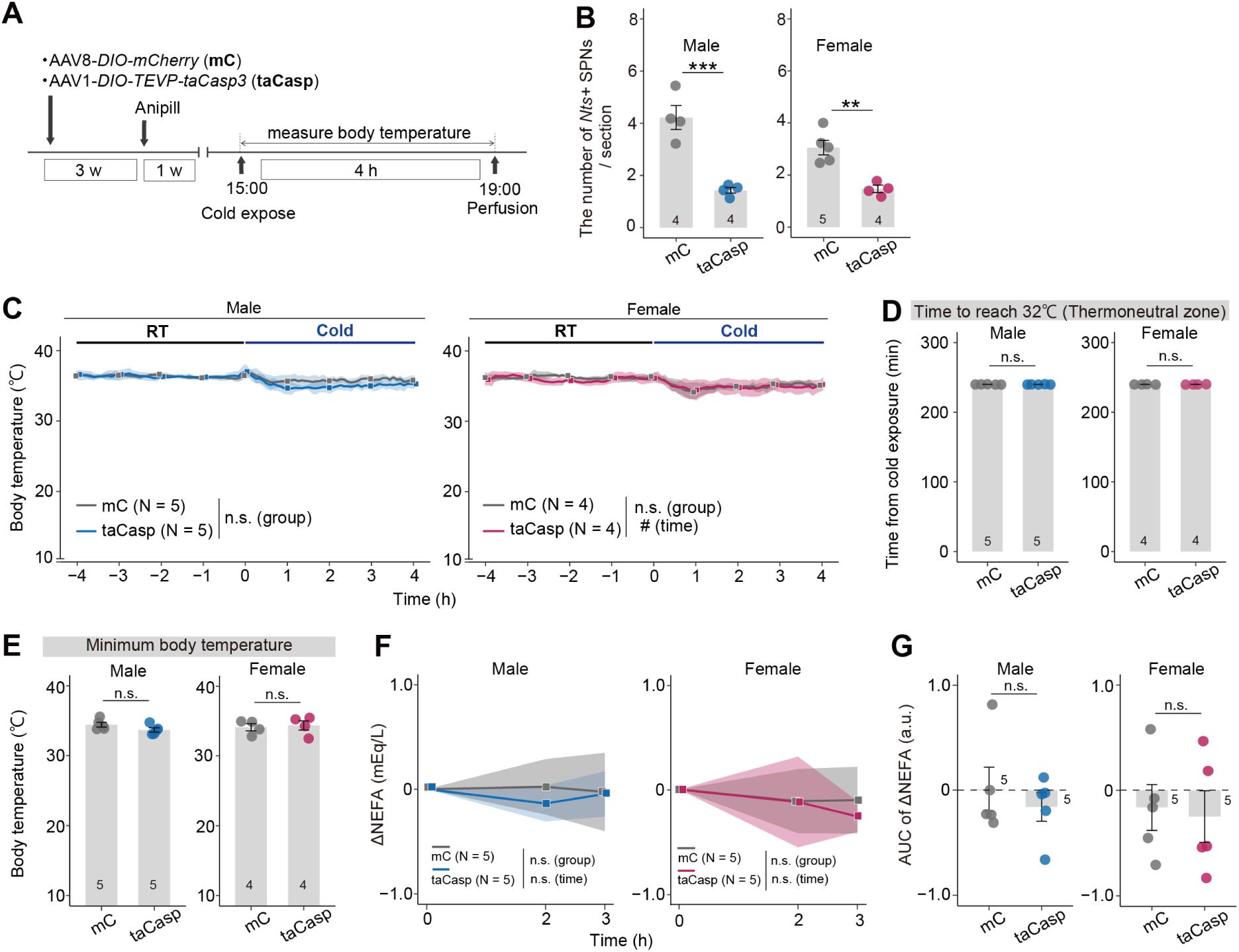
Thermal phenotype of *Nts*+ SPN ablation groups during cold exposure with ad libitum feeding, related to Figure 7. (A) Experimental timeline for measuring the Tc under cold exposure with ad libitum feeding. (B) Quantification of *Nts*+ SPN numbers per section in the control and taCasp groups. \*\**p* < 0.01, \*\*\**p* < 0.001 by two-sided Welch’s t-test. (C) Tc of mC (gray) and taCasp groups before and during cold exposure (4°C). Two-way repeated-measures ANOVA: group, time, and interaction effects, n.s. (male); group effect, n.s.; time effect, *p* < 0.01; interaction effect, n.s. (female). In females, the time effects at each time point relative to time 0 were further analyzed using one-way ANOVA followed by Bonferroni’s multiple comparison test (n.s.). (D) Latency to reach a Tc of 32°C in mC and taCasp mice. (E) Minimum Tc reached during cold exposure in mC and taCasp mice. For panels (D and E), no significant difference (n.s.) was observed using two-sided Welch’s t-test. (F) Changes in plasma NEFA levels (mEq/L) relative to the onset of cold exposure. Two-way repeated-measures ANOVA: group, time, and interaction effects, n.s. for both sexes. (G) AUC of ΔNEFA levels over 3 h of cold exposure. No significant differences (n.s.) in the Wilcoxon rank-sum test. The body temperature measurements in (C–E) and plasma NEFA measurements in (F, G) were performed in separate cohorts of mice. The number of animals in each group is shown. The error bars and shaded areas in panels (C) and (F) represent the standard deviations. The blue and pink dots represent male and female data, respectively.

**Table S1: Custom Xenium probe panel used in this study.**

This table lists the 100 genes selected for the custom Xenium add-on probe panel, together with their corresponding Ensembl gene identifiers and the number of probe sets assigned to each gene.

**Table S2: Metadata for the reanalyzed Blum et al. SPN single-nucleus RNA-seq dataset.**

This table contains the metadata used for reanalysis of the published SPN single-nucleus RNA-seq dataset from Blum *et al.* (2021), including the original sample identity (orig.ident), assigned cell type (celltype), and cell barcode (cell_id).

**Table S3: Numbers of spinal cord sections analyzed by Xenium.**

Rows are organized by sex, treatment condition, and animal ID, and columns indicate spinal segment groups from T1–4 through S2–4.

**Table S4: Metadata for Xenium-derived cholinergic neurons and SPNs.**

This table contains metadata for all cholinergic neurons detected in the Xenium dataset, including sample identity (orig.ident), assigned cholinergic neuronal subtype (Cholinergic_subtype), and unique cell identifier (cell_id).

